# Seven Millennia of Human Exploitation drove genomic Changes in Iberian Sheep

**DOI:** 10.64898/2026.02.03.702761

**Authors:** Pedro Morell Miranda, Juan Antonio Antolinos-Diaz, Paolo Mereu, Damla Kaptan, Marta Moreno-García, M. Ángeles Galindo-Pellicena, Monica Pirastru, Kıvılcım Başak Vural, Nicolò Columbano, Cristina Tejedor, Juan Luís Arsuaga, Giovanni G. Leoni, Amalia Pérez-Romero, Marta Francés-Negro, Eneko Iriarte, José Ignacio Royo Guillén, Salvatore Naitana, Ignacio de Gaspar, Manuel Rojo-Guerra, Mario Barbato, Colin Smith, José-Miguel Carretero, Mehmet Somel, Füsun Özer, Cristina Valdiosera, Torsten Günther

**Affiliations:** Human Evolution Program, Institute of Organismal Biology, Uppsala University, Uppsala, Sweden; Institute of Animal Systems Genomics, Faculty of Veterinary Medicine, Ludwig-Maximilians-Universität, Munich, Germany; Laboratorio de Evolución Humana, Universidad de Burgos, Burgos, Spain; Department of Biomedical Sciences, Università degli Studi di Sassari, Sassari, Italy; Department of Biological Sciences, Biology/Molecular Biology & Genetics, Middle Eastern Technical University, Ankara, Turkey; GI Arqueobiología, Instituto de Historia, CCHS-CSIC, Madrid, Spain; Centro Mixto UCM-ISCIII de Evolución y Comportamiento Humanos, Madrid, Spain; Departamento de Geología, Geografía y Medio Ambiente, Universidad de Alcalá de Henares, Spain; Department of Veterinary Sciences, Università degli Studi di Sassari, Sassari, Italy; Department of Prehistory and Archaeology, University of Valladolid, Valladolid, Spain; Department of Law and Humanities, European University of Madrid, Spain; Gobierno de Aragón, Zaragoza, Spain; Departamento de Anatomía y Embriología. Facultad de Veterinaria. Universidad Complutense de Madrid, Madrid, Spain; Department of Veterinary Sciences, Università degli Studi di Messina, Messina, Italy; Archaeology and History department, La Trobe University, Melbourne, Australia; Faculty of Letters, Department of Anthropology, Department of Social Anthropology, Hacettepe University, Ankara, Turkey; Centro Nacional de Investigación sobre Evolución Humana. Burgos, España; Science for Life Laboratory, Institute of Organismal Biology, Uppsala University, Uppsala, Sweden; Center for Paleogenomics, Stockholm University, Stockholm, Sweden

**Keywords:** Sheep, Ancient DNA, Demography, Wool, Genomics

## Abstract

As one of the first domestic livestock species, sheep have played a fundamental role in human societies since the Neolithic. However, their demographic history remains poorly understood. To shed light on the demographic dynamics of sheep at the western edge of the Mediterranean, we sequenced 22 ancient sheep genomes from Iberia (dating from 7, 270 to 1, 615 calBP, sequenced up to 8.74 × coverage) along with one modern European mouflon from Corsica. We provide evidence for an initial maritime introduction into Iberia, and show that European mouflons are descendants of feralized Neolithic sheep. Furthermore, we identify a secondary influx of “Eastern” genetic ancestry coinciding with the arrival of human Steppe ancestry in Iberia – an event that likely aligns with the spread of woolly sheep across Europe. A third population expansion is observed during the Roman period, a time when historical sources reference the trade of fine-wool sheep. This Roman-era expansion appears to have significantly influenced the genetic makeup of modern European sheep, contributing to the development of popular modern-day breeds such as Merino. In addition to these major events, we see indications of additional, minor episodes of prehistoric gene flow into the Iberian population, suggesting that western European sheep experienced more dynamic demographic changes than other domestic animals, humans, or sheep populations elsewhere. Together, these results highlight the dynamic history of Iberian sheep populations and demonstrate how human cultural and demographic shifts have left their hoofprints in the sheep gene pool, marking them as a valuable proxy for understanding the human past.

## Introduction

Sheep (*Ovis aries*) have been a key resource in human societies since they became first domesticated in the North-Western Fertile Crescent about 10000 − 12000 years ago (1– 6), along with playing an important role on most societies due to their symbolic and cultural significance. From these early domestic herds, they expanded both eastward and westward, accompanying human populations during the Neolithic Expansion (7). This expansion followed different and complex routes along different regions in Eurasia (8), with two main migration axes into western Europe: an expansion following the Mediterranean islands and coastline, associated with the arrival of the *Cardial* and *Impressa* ceramics to southern Europe, and the Danube-Rhine axis, associated with the LBK (*Linnearbandkeramik*) culture that spread over Central and Northern Europe. It was the coastal Mediterranean route that brought farmers as well as livestock and crops to Iberia, where sheep and goats rapidly became important food sources (9).

Following this initial expansion of “hairy” sheep, phenotypically similar to modern-day mouflons, at least one secondary migratory wave has been assumed to have occurred before the Bronze Age, associated with the arrival of the “woolly” phenotype to Europe (10, 11). This migration also occurred during a general shift in the way domestic animals were exploited by early farmers. From breeding them for the production of primary resources, such as meat and hides, they transitioned to focus on the production of more sustainable products, such as milk, wool, fertilizer or traction (12, 13). However, great uncertainties about when this expansion took place and from where it originated remain, as wool is rarely preserved in the archaeological record and most evidence for the aforementioned change has been inferred from differences in culling patterns and the appearance of wool-processing tools in the archaeological record (10, 11, 13). Due to the limited availability of early ancient wool, several genetic studies have used DNA information from modern sheep breeds to trace this secondary expansion. Patterns seen in endogenous polymorphic retroviruses (14), the Y chromosome (15), and low-density genotype data (16) independently suggest additional “Eastern” gene flow into Europe after the initial domestication and expansion of sheep without providing an exact timing for when that happened. We want to stress that in this case, “Eastern” means that it has a higher affinity with modern sheep from outside of Europe and not any concrete part of Asia.

There are, however, limitations to the information that genomic data from modern breeds can offer us about their past, as many patterns have been erased or obscured by later events. Archaeogenomics provides the possibility to directly study past populations through ancient DNA (aDNA). In pigs, aDNA showed that the close relatedness of modern European pigs with European wild boars was caused by massive gene flow from wild to domestic populations and not an independent domestication outside of the Fertile Crescent (17). Similarly, studies with ancient horses suggest that modern domestic horses do not descend from the initial domestication carried out by the Eneolithic Botai culture, but from a later event inferred to have occurred in the Volga-Don region (18–20). Other domestic species’ demographic history research, like dogs (21, 22), cattle (23), and goats (24, 25), also have benefited from the increased resolution offered by aDNA to disentangle how they were domesticated and/or how they expanded and evolved since. In sheep, existing research focused on single geographic regions, and none included a dedicated multi-period temporal sampling that would allow the detection of population transitions. Other than matrilineal and patrilineal studies on a local scale (26, 27), several publications exist that analyzed the demographic histories of sheep at a local level. These studies focused on Neolithic Anatolia (28, 29), Neolithic and Bronze Age Kyrgyzstan (30), Iranian Antiquity (31) and the Late Neolithic and Middle Ages from Northern Europe (32). A recently published study (6) with a wider regional and temporal sampling described how the early domestic sheep populations were formed and how they are related to Northwestern European flocks from later time periods.

Despite the growing availability of aDNA data, large regions of Eurasia remain unsampled. Further, no study has yet conducted a comprehensive time-transect from the same region to detect population changes over time. Iberia and South-Western Europe are one of the blank spots where no temporal record exists of the sheep gene pool, despite the historical and current importance of the species and the presence of archaeological sites with good DNA preservation, as known from e.g. humans (33–35), cattle (36) and horses (19, 37). At the Atlantic coast of Europe, Iberia represents the western end of sheep expansion and the putative origin of some historically important breeds like the fine-wool Merino (38). To investigate the evolution and demographic history of Iberian sheep, we sequenced the genomes of 22 ancient sheep remains with coverage values ranging from 0.0022 to 8.74 × dating from the Neolithic to the Roman Age from two cave sites in Northern Iberia: the Neolithic site of Els Trocs in the Spanish Pyrenees, as well as El Portalón, a site with a detailed stratigraphic record located in the Atapuerca complex that spans from the Neolithic to the Visigothic periods (Figure 1A and B, Table 1, Supplementary Table 1). Additionally, we generated a medium coverage genome (8*×*) for one modern European mouflon from Corsica (*Ovis aries musimon*) (Supplementary Table 2), a population considered to represent feral Neolithic sheep with limited introgression from later domestic breeds (39, 40). We compared them with modern and ancient genomic data from other locations (15, 28, 29, 32) (Supplementary Tables 1 and 2). This temporal dataset allowed us to create a chronology of the demographic changes in sheep populations in Iberia, illustrating how the gene pool changed in connection with human cultural and demographic shifts.

**Table 1.** Summary information of the novel ancient samples included in the population genomic analyses for this study:

| Sample | Site | Archaeological association | Dating (cal BP) | Sex | Coverage | MT Haplogroup |
| --- | --- | --- | --- | --- | --- | --- |
| APOR-013 | El Portalón | Roman | 1819 – 1615 | XY | 0.97× | B1a2 |
| APOR-002 | El Portalón | Bronze Age | 3826 – 3580 | XX | 8.74× | B1a1 |
| APOR-003 | El Portalón | Bronze Age | 3878 – 3702* | XX | 2.47× | B1a2a1 |
| APOR-004 | El Portalón | Bronze Age | 4061 – 3727 | XX | 5.92× | B1a |
| APOR-014 | El Portalón | Bronze Age | 3878 – 3702* | XX | 0.05× | B1a1 |
| APOR-015 | El Portalón | Chalcolithic | 4843 – 4623 | XX | 0.10× | B1a1 |
| APOR-016 | El Portalón | Chalcolithic | 4811 – 4529 | XX | 0.11× | B1a1 |
| APOR-005 | El Portalón | Chalcolithic | 5041 – 4851 | XX | 2.66× | B1a1b |
| APOR-011 | El Portalón | Chalcolithic | 5587 – 5331 | XY | 3.27× | B1a2a |
| APOR-008 | El Portalón | Neolithic | 7270 – 7024 | XX | 1.32× | B1a1 |
| APOR-009 | El Portalón | Neolithic | 7260 – 6992* | XY | 8.37× | B1a1b |
| ATRO-001 | Els Trocs | Neolithic | 4571 – 4406* | XX | 0.06× | D,D_HM236181 |
| ATRO-002 | Els Trocs | Neolithic | 4571 – 4406* | XY | 0.36× | B1a |
| ATRO-003 | Els Trocs | Neolithic | 6645 – 6412* | XX | 0.04× | B1a2a1 |
| ATRO-004 | Els Trocs | Neolithic | 7159 – 6893* | XX | 0.16× | B1a2a1 |
| ATRO-007 | Els Trocs | Neolithic | 7159 – 6893* | XX | 0.03× | B1a2a |
| ATRO-008 | Els Trocs | Neolithic | 7159 – 6893* | XX | 0.03× | B1a |

\* Contextual dates from other radiocarbon dating of organic material found in the same stratigraphic layer. The Els Trocs samples show the whole range of dates from their corresponding layer.

**Fig. 1.**
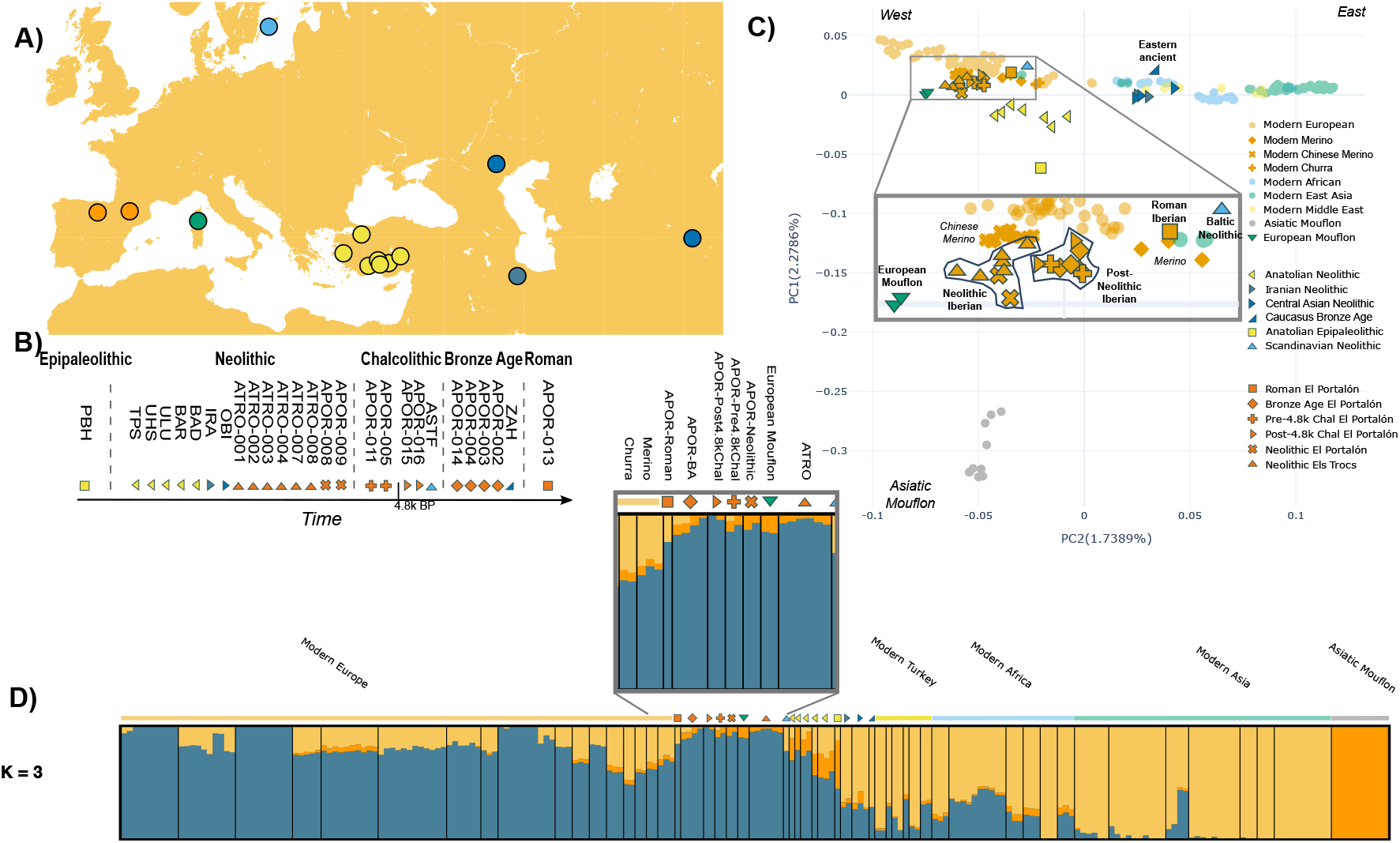
(A) Geographic distribution of the investigated archaeological sites, plus the sampling site for the modern Corsican Mouflon and the ancient individuals used as reference. The map was generated using the Open Source software pyGMT (41). (B) Chronological representation of the ancient samples under study. The timeline is not to scale. Period classifications follow their regional chronology, which means that dates of the samples may not correlate with sample ages across geographical regions. (C) PCA plot. Modern individuals were used to calculate the components. Ancient samples were projected and are highlighted by saturated colors and dark margins. The European mouflons, which were not projected, have also been highlighted for clarity. (D) ADMIXTURE plot with K=3. Ancient Iberians are zoomed in to facilitate the comparison with modern Iberian sheep (Merino and Churra). All highlighted populations follow the color schemes and symbols described in the PCA legend.

## Results and Discussion

Ancient DNA was extracted from teeth and attached jaw bones from 23 samples, from which 22 produced enough DNA to continue our pipeline. Prepared libraries were shotgun sequenced on Illumina NovaSeq 6000 machines. The generated sequence data shows fragmentation and postmortem deamination as expected for authentic endogenous aDNA (See Supplementary Figure 1). Eight samples were directly radiocarbon dated to between 7270 − 7024 and 1819 − 1615 cal BP. Based on the number of reads mapping to the X chromosome versus the ones mapping to the autosomes, we observe a significant abundance of females at our sites, where 17 individuals were identified as female and 4 individuals as male (*p*-value= 0.0072). This pattern is similar to the one seen in a previous study in Anatolia (29), which may be related to the exploitation pattern (i.e. culling the young males). Importantly, most of the samples from El Portalón belong to layers of no human occupation, but active penning of sheep at the site, probably during winter. This fact, added to the seasonality, may bias the ratio at which female versus male carcasses were deposited at the site. As expected, most samples from ancient Iberia belong to MT haplogroup B (Table 1), the most common haplogroup in modern European sheep, which has been associated with the initial western expansion (26, 28). The only exception is a Neolithic sample from Els Trocs (ATRO-001), which belonged to haplogroup D, making this the only observation of haplogroup D in an ancient European sheep.

For the downstream analyses, samples with coverage below 0.01 × were discarded (3 samples from El Portalón and 2 samples from Els Trocs, described in Supplementary Table 1), leaving 17 samples for population genomic analysis (Table 1). We describe the results of the genomic analysis in chronological order below. Throughout this study, our terminology follows the Iberian chronology.

### Expansion of Neolithic sheep into Europe

Sheep first arrived in Iberia during the Neolithic after following the coastal Mediterranean route. An exploratory Principal Components Analysis (PCA) (Figure 1C) shows a gradient between modern wild Asiatic mouflons and modern domestic sheep on PC1, with the ancient individuals mostly aligning with chronology. The Epipaleolithic Anatolian is placed between the wild Asiatic mouflons from Iran and modern domestic sheep, and the Early Neolithic samples from Anatolia are intermediate between this sample and more recent sheep. The only exception to this pattern is the modern European mouflons, which show values similar to those of the Neolithic Iberian samples on PC1 but are also drifted apart from them along PC2 towards the Northwestern European breeds. This position supports the hypothesis that European Mouflons are feralized Neolithic Mediterranean sheep. PC2 generally captures the West-to-East geographical structure in modern sheep breeds as described in previous studies (16, 39, 42, 43). Like all Western Eurasian sheep, Neolithic Iberian sheep also display a shift towards lower values than earlier Neolithic Anatolian populations along PC2, while later Iberian sheep display higher values, although never leaving the space covered by modern European breeds. This pattern is consistent with the results of the ADMIXTURE analysis (Figure 1D) displaying a distinctive lack of an Eastern component (in yellow) during the Neolithic that appears in later Iberian sheep and is common nowadays in Mediterranean and other European sheep. Neolithic Iberian sheep also display a low amount of the component associated with wild Asiatic mouflons, which correlates with PC1 and decreases in more recent samples from Iberia, almost disappearing in modern sheep.

These signals are corroborated by reconstructed admixture graphs, aiming to describe the demographic history of Iberian sheep. The two methods used for graph reconstruction, qpGraph (44) and OrientAGraph (45), show a similar pattern, highlighting that the European mouflon diverged from the initial domestic Neolithic sheep expanding through the Mediterranean (Figure 2A, Supplementary Figure 5 A-G). The Neolithic sheep from Iberia and the European Mouflon share the same basal ancestry from Anatolian Neolithic sheep and Asiatic Mouflons (similar to PC1 and ADMIXTURE), consistent with them having a single origin and presenting somewhat more primitive sheep breeds compared to later groups. To corroborate this result, we performed *f*_4_ statistics in the shape of *f*_4_(*Urial, EpipaleolithicAnatolian*; *X, APOR* − *N eo*) and *f*_4_(*Urial, X*; *CaucasusBronzeAge, APOR* − *N eo*). In the first test (Supplementary Figure 4A), European Mouflons are not significantly different from 0, similar to the ancient populations, while in the second (Supplementary Figure 4B), they are the only non-prehistoric population that is significantly drifted towards the Iberian Neolithic sheep, which also supports their feral status. The Neolithic Iberian sheep also show lower genetic diversity than their Anatolian source population (Figure 2C, Supplementary Table 7), measured as the conditional nucleotide diversity (46). This could be expected since the Mediterranean expansion, which may have partly involved transport on boats, was likely associated with a series of bottlenecks on a “stepping-stone” migration. Notably, we see a slight increase in genetic diversity between the North-Eastern Iberian Neolithic Els Trocs and Neolithic El Portalón in central Northern Iberia. Considering the locations and chronologies of the two sites – Els Trocs being closer to the putative point of entry of the expansion into Iberia – this is consistent with the arrival of a small stock of Neolithic sheep to North-Eastern Iberia, which then slowly increased in population size, recovering from the migrationassociated bottleneck. This could also be supported by the Neolithic faunal record of El Portalón, where caprine representation increases considerably from Early Neolithic to Late Neolithic (47).

**Fig. 2.**
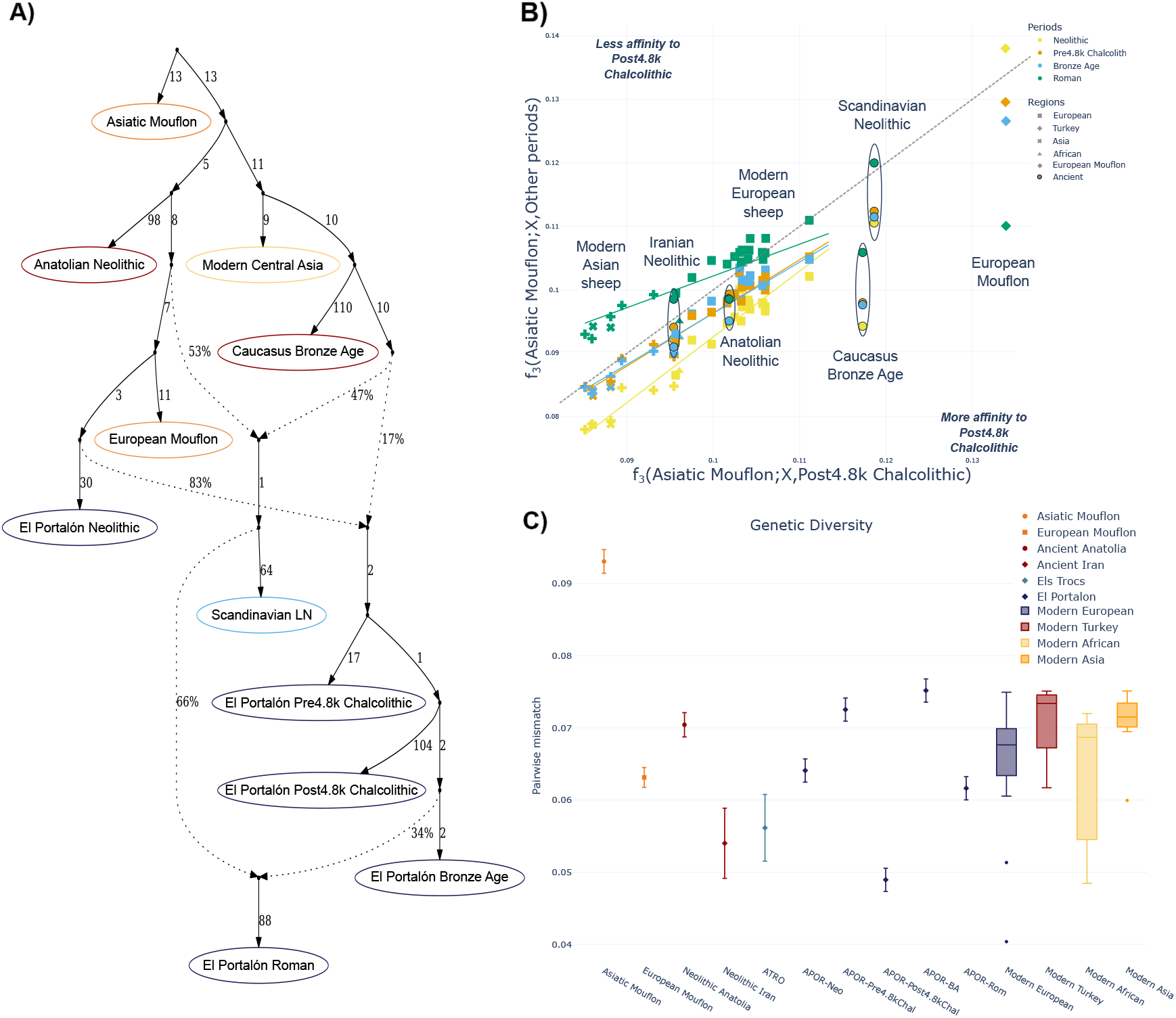
(A) Admixture Graph for ancient sheep inferred with qpGraph assuming 3 admixture events. Other admixture graphs are shown in Supplementary Figure 5. (B) Comparison between the results of Outgroup-*f*_3_ statistics for the Post-4.8k Chalcolithic samples and samples from other periods in El Portalón, in the format *f*_3_ (*AsiaticM ouf lon*; *X, ElP ortal*ó*n*). Full commaprisons of all periods are shown in Supplementary Figure 3. (C) Pairwise conditional molecular diversity. Mouflons and ancient groups are represented by dots, while modern populations are aggregated by region. Ancient groups are also ordered chronologically.

### Increased Eastern Ancestry during the Chalcolithic

Compared to the preceding Neolithic individuals, Chalcolithic Iberian sheep (dated between ∼ 5,400 and ∼ 4,400 cal BP) display a slight shift towards modern groups in the PCA (Figure 1C). The ADMIXTURE results even suggest a small difference between the two earlier Chalcolithic samples and the later samples, dated to after ∼ 4,800 cal BP, which show more modern European ancestry and have a lower level of Asiatic mouflon component (Figure 1D). In order to explore this genetic pattern further, we divided the Chalcolithic samples into Pre-4.8k and Post-4.8k Chalcolithic, as this transition predates the cultural Pre-Bell Beaker and Bell Beaker classification traditionally used in this region (49, 50). The admixture graphs attribute the slight difference observed in the ADMIXTURE results to external gene flow: OrientAGraph infers more basal ancestry in Pre-4.8k sheep, suggesting another gene flow event into Iberia (Supplementary Figure 5). qpGraph summarizes all Chalcolithic groups as a mixture of 83% Iberian Neolithic ancestry and 17% ancestry related to a Bronze Age individual from Kalmykia, close to the Caucasus (Figure 2A), confirming at least one gene flow event from non-Iberian sources around during the Chalcolithic. These results are corroborated by shared drift with modern breeds inferred through Outgroup-*f*_3_ statistics (44): compared to Neolithic sheep, Pre-4.8k Chalcolithic sheep show an elevated level of genetic similarity with all (not only European) modern sheep breeds (Supplementary Figure 3). Notably, this pattern extends to Eastern ancient sheep from Neolithic Iran and Bronze Age Caucasus, suggesting some level of increased “Eastern” ancestry (i.e., non-European) during the Pre-4.8k Chalcolithic, as also inferred by qpGraph (Figure 2A). This observation is in line with a subtle increase in Eastern ancestry observed in Chalcolithic Eastern Europe (6). The exact geographic origin of this gene flow was unclear, but it pre-dates any Steppe-related expansions (6) and may have extended all the way to Western Europe, as suggested by the similar signal in the Iberian Pre-4.8k Chalcolithic.

Later during the Chalcolithic, we observe another increase in the affinity of Post-4.8k Chalcolithic Iberian sheep to all modern breeds, when compared to either Neolithic, Pre-4.8k Chalcolithic, or even Bronze Age samples (Figure 2B). The Roman Age sample is the only exception that displays higher levels of shared drift with modern breeds, which was expected for temporally later populations. Both Post-4.8k Chalcolithic and Roman sheep display significantly higher levels of affinity with modern breeds in general, and Eastern breeds in particular, than populations from earlier periods (Figure 2B, Supplementary Figure 3). The Post-4.8k Chalcolithic samples also show an outstanding level of shared drift with the Bronze Age Caucasus sample and the Late Neolithic sample from Gotland, in contrast to the Pre-4.8k Chalcolithic sheep, which displayed similar levels of affinity as the other groups across comparisons (Supplementary Figure 3). Both of these samples have been shown to contain large proportions of ancestry originating from the Pontic-Caspian Steppe (29), suggesting an influx of such ancestry into Iberia during the Post-4.8k Chalcolithic. Notably, the Eastern ancestry in Iberian sheep as well as Late Neolithic Scandinavian sheep are inferred to originate from a similar source by qpGraph (Figure 2A). These results, in combination with a substantial drop in genetic diversity compared to previous and later ancient sheep (Figure 2C), suggest that Post-4.8k Chalcolithic sheep belonged to a different stock that arrived recently to Iberia and received its Eastern Ancestry from a different route and source than the Pre-4.8k Chalcolithic samples.

To further explore this Eastern affinity, we calculated individual-based *f*_4_ statistics using Altay sheep, a modern breed from Western China, as a proxy for Eastern ancestry. We used the oldest directly dated Neolithic sample in our dataset (APOR-008; 7270 − 7024 cal BP) as a baseline for the initial Iberian sheep ancestry. While Eastern Eurasian sheep are symmetrically related to the Pre-4.8k Chalcolithic and Neolithic sheep, we see a significant increase in Eastern ancestry in our Post-4.8k Chalcolithic sheep (Figure 3A; statistically significant for APOR-015). This pattern is also replicated when we use other Eastern references, such as Duolang, another Central/East Asian sheep breed (Supplementary Figure 6A), but also archaic Northern European breeds, such as East Friesian, Gotland or Drenthe Heathen (Supplementary Figure 7). Interestingly, the Pre-4.8k Chalcolithic samples also show a significantly increased amount of Eastern ancestry when the source of the Eastern Ancestry is a Southwestern Asian sheep breed such as Afshari or Awassi (Supplementary Figure 6B, C), which might indicate their generally increased Eastern affinity compared to the Neolithic, as described in previous literature on the Balkans (6).

**Fig. 3.**
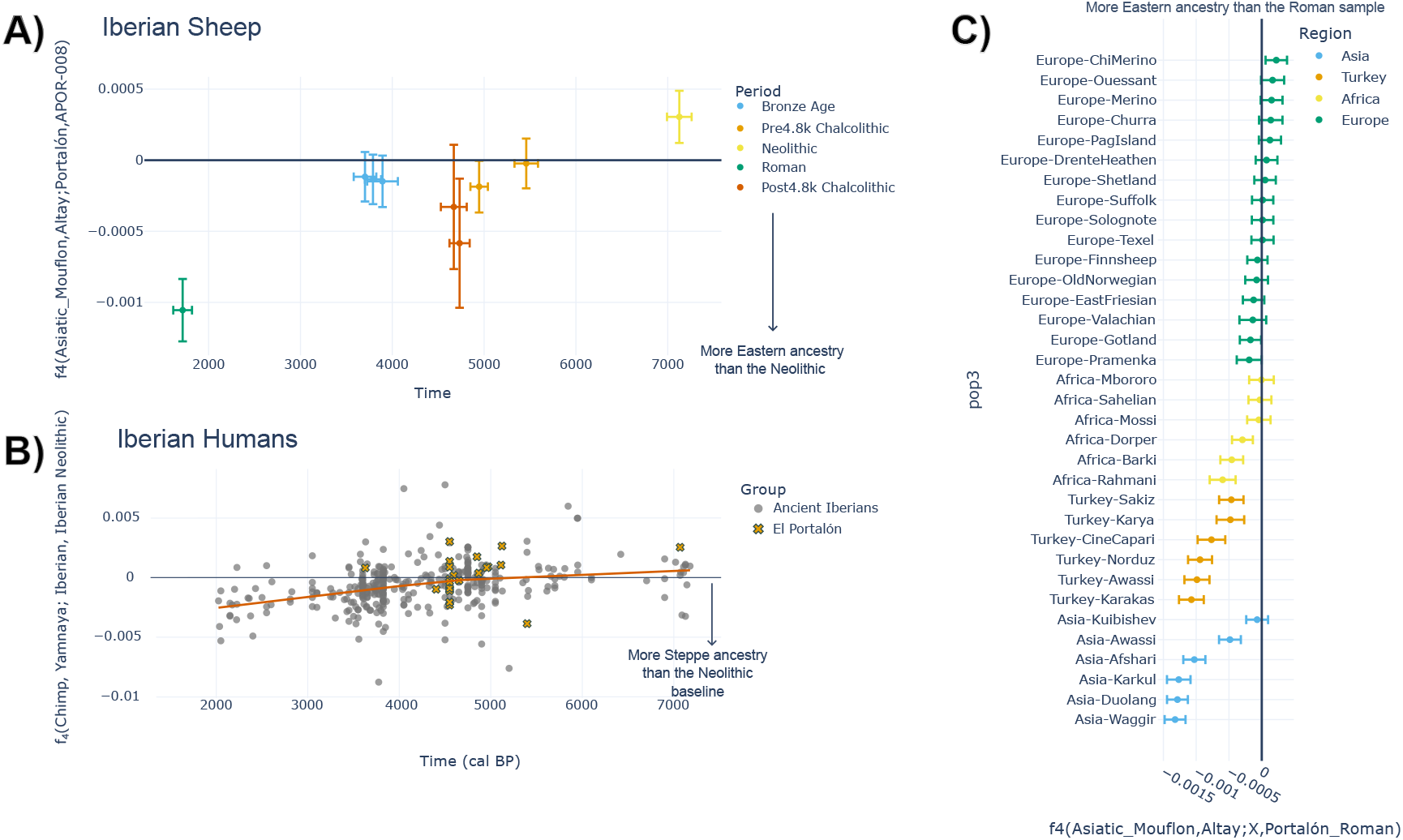
(A) Temporal change of the affinity to Eastern sheep polarized between the oldest Neolithic sheep from El Portalón and modern Altay sheep in the form *f*_4_ (*Asiatic Mouflon, Altay*; *ElP ortal*ó*n, ElP ortal*ó*nN eolithic*). (B) Temporal change of the affinity to Steppe ancestry in Iberian humans, polarized between the oldest Neolithic sample and ancient Yamnaya samples from Eastern Europe in the form *f*_4_ (*Chimp, Y amnaya*; *Iberian, IberianNeolithic*). The dataset is a subset of all Iberian samples from the Allen Ancient DNA Resource (AADR) V54.1.p1 (48). (C) Genetic affinity of modern breeds to Eastern ancestry polarized between the Roman sample and modern Altay sheep in the form *f*_4_ (*Asiatic Mouflon, Altay*; *M odern, APOR* − *Roman*).

Similar to previous studies (6), we cannot link the Pre-4.8k Chalcolithic influx of low levels of Eastern ancestry to known prehistoric events. However, the significant increase in Eastern sheep ancestry during the Post-4.8k Chalcolithic coincides with the arrival of human Steppe ancestry in Iberia (35) (Figure 3B), with the first individual dated to 4843 − 4623 cal BP. The first human individual with Steppe ancestry from the neighboring site of El Mirador, less than a kilometer from El Portalón, dated to 4825 − 4441 cal BP (51). This suggests that the changes in genomic ancestry in the two species were directly or indirectly connected. Furthermore, one may speculate that such a broad movement, which also changed the sheep gene pool in other parts of Europe (6, 29, 32), was associated with a transition in the way sheep were exploited, such as an increased use of their fibers. While the earliest direct archaeological evidence of wool processing in Iberia dates to the Bronze Age or early Iron Age (52, 53), these genetic changes overlap also with an increase in secondary exploitation of sheep in Northern Iberia (54) and, therefore, the potential time for the spread of woolly sheep (1, 26, 38). It should also be noted that in El Portalón and other sites, artifacts interpretable as spindle whorls are present from the Neolithic to the Bronze Age (55). The Iberian Chalcolithic sheep in this study probably represent the descendants of a complex process of secondary westward expansions during the Chalcolithic, whose genetic signature can still be detected in some traditional European sheep breeds today. These breeds, such as Drenthe Heath, Gotland or East Friesian, are characterized by retaining coarse straight hair in addition to long, usually dark wool, and some also exhibiting seasonal molting (56). These are archaic characteristics modern woolly breeds have mostly lost. While previous studies have assumed these breeds were, as the European Mouflons, a remnant of the first sheep expansion into Europe (14), the genetic connection between extant coarse wool sheep and Chalcolithic sheep supports this secondary expansion into Europe as the initial expansion of archaic woolly sheep westwards.

### Admixture during the Bronze Age

The El Portalón samples from the Bronze Age (dated to after ∼ 3,900 cal BP) show certain genetic similarities to the Chalcolithic Iberian sheep. While both Chalcolithic groups and the Bronze Age samples show similar values in both PCA and ADMIXTURE (Figure 1C and D), we see important differences between Bronze Age and Chalcolithic sheep in their genetic affinity with modern populations (Figure 2B). In contrast to the expectations from chronology, the Bronze Age samples share less genetic drift with most modern breeds than the Post-4.8k Chalcolithic samples (Figure 2B). Interestingly, there is little difference between the values of the Pre-4.8k Chalcolithic and Bronze Age sheep in the Outgroup-*f*_3_ comparison (Figure 2B, Supplementary Figure 3). We also see that these sheep follow the pattern of increasing genetic diversity seen in previous periods, with the exception of the Post-4.8k Chalcolithic individuals (Figure 2C).

Furthermore, the admixture graph inferred using qpGraph places the Bronze Age sheep from El Portalón as descendants of the same branch the Chalcolithic populations belonged to (Figure 2A), supporting the idea that this population is mostly continuous with previous domestic stock from the region. On the other hand, the reconstruction of the admixture graphs with OrientAGraph (Supplementary Figure 5A-F) consistently places the Bronze Age samples as an outgroup to earlier domestics from Iberia, independently of the number of migration edges. These results could be explained as the local Pre-4.8k Chalcolithic population mixing with the new stock that came in during the Post-4.8k Chalcolithic, and diffusing the incoming ancestry into the Iberian gene pool over time. The earlier group of sheep could represent a local stock that was adapted to local Iberian environments, while the second population carried genetic variants for new and desirable production traits (i.e. wool and maybe milk). Alternatively, different types of sheep stock may have lived in parallel and in isolation on the Iberian Peninsula.

Nowadays, Iberian herders follow seasonal routes of sometimes hundreds of kilometers in a process known as transhumance, which can be traced back at least to the Roman period and probably even earlier (57), as isotopic evidence suggests seasonal vertical mobility of sheep herds in early Neolithic sites such as Els Trocs (58). Such seasonal migratory routes could have facilitated both scenarios, either by providing opportunities for the mixing of groups or the isolation of populations within Iberia due to climate and cultural/political barriers, leading to changing routes.

Zooarchaeological analyses of sheep and goat remains from El Portalón suggest that there was an increased exploitation of secondary products during the Chalcolithic. However, this exploitation decreased during the Bronze Age at this and other sites of the North and East of Iberia, with two-thirds of the exploitation reverting to a focus on meat production (59, 60). This intensification in the exploitation of secondary products can also be observed in the lipid analyses of ceramic vessels from El Portalón. Ruminant dairy lipids are almost absent during the Neolithic, followed by a progressive increase during the Chalcolithic and Bronze Age (47). Combined, these results suggest herders in Northern Iberia developed their breeding practice to combine systematic secondary production like milk and wool from mature females with high meat productivity. This pattern of secondary exploitation is reversed in the Argaric culture of Southern Iberia (61), supporting the idea of different practices existing in parallel on the Iberian Peninsula and, consequently, cultural or political barriers shaping the Iberian sheep gene pool. Finally, we also need to consider that the genetic differences in the Bronze Age population could also be connected to an additional influx of sheep from outside Iberia (as suggested by more complex admixture graph models; Supplementary Figure 5), which is difficult to pinpoint due to the relatively minor genetic change and the lack of good candidate populations for the source of such gene flow.

### Further introductions during the Iron Age or Roman period

After the Bronze Age, our dataset has a sampling gap of about two millennia. We observe an evident genetic discontinuity between our earlier Iberian sheep and the single Roman period individual (APOR-013, dated to the early 3*rd* century CE; Table 1) analyzed in this study. Compared to preceding time periods, PCA and ADMIXTURE show this sample more similar to modern Eastern Eurasian sheep (represented by lower values in PC2 in Figure 1C and by the yellow component in Figure 1D) with almost no component associated with the wild Asiatic mouflons still present in older Iberian sheep (Figure 1D). In fact, this individual shows a pattern more similar to modern Iberian breeds like Merino or Churra than to any of the other ancient sheep.

This individual is also sharing a significantly higher level of genetic drift with all modern breeds than older Iberian sheep, particularly with modern European long-tailed breeds and breeds from Asia and Africa (Figure 2B and Supplementary Figure 3). Our qpGraph admixture graph corroborates these results and also points to an explanation: the Roman individual is modeled as a mix of the El Portalón Bronze Age sheep (34%) and a foreign population with enriched Eastern Ancestry (66%). This population, distantly related to the Scandinavian Late Neolithic sheep, was also a mix of basal Anatolian Neolithic and Eastern Neolithic ancestry, on the graph represented by a Caucasus Bronze Age sheep (4099 − 3945 Cal BP) from Kalmykia, Russia. All these results highlight the different genetic background of this individual, suggesting an intentional importation of sheep with improved wool from the Eastern Mediterranean or from regions outside the Roman Empire (e.g., Tarentum and Colchis, in modern Italy and Georgia, respectively) during this period, as suggested by historical Roman sources (38). These sources were interpreted as describing the origin of fine-wool sheep, and in particular as the origin of the Merino breed that became extremely popular during the Middle Ages in Iberia and later other parts of Europe. However, our results show a more widespread pattern of mixing of this new ancestry with local sheep stock in Iberia. The Roman population also seemed to have a lower genetic diversity compared to the Bronze Age. As this population seems to be heavily admixed, this may suggest a change in breeding practices during this period, with stronger selective pressures and more intense breeding. This value, however, should be interpreted with caution, as the pairwise comparison was performed using one of the lowest-quality genomes sequences in this study (APOR-001), and was calculated on only 2, 531 sites, which is several orders of magnitude fewer than in other comparisons (Supplementary Table 7).

As the Roman sample was the ancient individual with the highest level of shared drift with modern Eastern Eurasian sheep (Figure 3A), we also used *f*_4_ statistics in the form of *f*_4_(*Asiatic Mouflon, Altay*; *M odern, APOR* − *Roman*) to explore whether any modern sheep breed carries more Eastern ancestry than the Roman individual (Figure 3C). While non-European sheep display an excess of allele sharing with Eastern Eurasian sheep, most European breeds do not, with the exception of some Eastern Mediterranean and Northern European breeds. The fact that Mediterranean and Western European breeds show similar levels of Eastern ancestry as the Roman sample suggests that there has been no significant new gene flow from outside of Europe into South-Western European stock since the Roman period.

## Conclusions

Our temporal dataset allowed us to trace more than seven millennia of sheep evolution in the Western Mediterranean. While our results suggest the isolation of the European mouflon since the Neolithic, this is hardly the case for managed Iberian sheep, whose herds have received at least three additional separate waves of gene flow from Eastern populations after the original expansion. The ancestry shift during the Chalcolithic seems to have consisted of two migrations from the East into Europe with different sources, followed by a potential additional wave during the Bronze Age.

The final major ancestry shift predates or coincides with the Roman occupation of Iberia, a period that reshaped European cultures in many ways, and our results suggest that they might have had a profound impact on the gene pool of European sheep breeds today, laying the foundation for modern production sheep breeds. While some authors have hypothesized that Roman translocations were the origin of fine wool sheep breeds such as the Merino (38), our results suggest a wider influence in most modern European breeds.

As a primarily genetic study focused on a single geographic region, our work lacks direct evidence of wool processing at the investigated sites and genomic data from potential source populations. Therefore, we can only correlate changes in the Iberian sheep gene pool with wool production or quality, without demonstrating a causal link. While future studies may address these gaps with additional data, it is plausible that shifts in the sheep gene pool reflect the preferences of prehistoric and historic farmers and consumers for improved livestock. The temporal genetic groups show similarities to modern groups with different coat and wool types, making wool the most likely trait associated with these transitions.

Our results highlight the dynamic nature of sheep herding in the Western Mediterranean and the major influence human cultural changes and movements had on sheep evolution in Europe, particularly in Iberia. From what we know so far, the Iberian sheep gene pool appears to have been more dynamic than that of humans or other domestic animals within the same time frame. Western Europe also appears to be unique in this case, as other regions such as Northern Europe and large parts of Asia (6) show continuity in the sheep gene pool deep into prehistory. The demographic changes in Iberian sheep had a profound impact on the morphology and gene pool of modern landraces and commercial breeds alike. Furthering our understanding of these processes is key to comprehending how these breeds developed and evolved into the stocks we can find today.

## ACKNOWLEDGEMENTS

We are incredibly grateful to the excavation teams at the two sites included in this study, Els Trocs and El Portalón de Cueva Mayor. The Atapuerca research project is financed by the Spanish Ministerio de Ciencia, Innovación y Universidades project nº PID2024-156477NB C-33. The field work at the Sierra de Atapuerca sites is funded by the Junta de Castilla y León and Fundación Atapuerca. This work was supported by grants from the Swedish Research Council Vetenskapsrådet (2017–05267) and Erik Philip-Sörensens stiftelse (G2020-035) to TG; an Ax:son Johnson grant (F20-0274) to PMM; a Ramón y Cajal (RYC2018-025223-I) from the Ministerio de Ciencia, Innovación y Universidades, Agencia Estatal de Investigación, AEI to CV and a Beatriz Galindo Fellowship (BGS220-461AA-69201) from the Ministerio de Ciencia, Innovación y Universidades (MICIU) to CS. MG-P is supported by a postdoctoral research contract within the PHS-2024/PH-HUM-469 programme of the Comunidad de Madrid. Processing of aDNA was performed by the SciLifeLab Ancient DNA Unit. Sequencing was performed by the SNP&SEQ Technology Platform in Uppsala, part of the National Genomics Infrastructure (NGI), Sweden and Science for Life Laboratory. The SNP&SEQ Platform is also supported by the Swedish Research Council and the Knut and Alice Wallenberg Foundation. The computations and data handling were enabled by resources in projects SNIC 2020/15-300, SNIC 2021/22-763, SNIC 2022/22-949, SNIC 2022/22-985, UPPMAX 2023/2-31, NAISS 2023/2-19, NAISS 2023/22-1053, NAISS 2024/22-851, NAISS 2024/22-1281, UPPMAX 2025/2-14 and UPPMAX 2025/2-93, provided by the National Academic Infrastructure for Supercomputing in Sweden (NAISS) and the Swedish National Infrastructure for Computing (SNIC) at Uppmax, partially funded by the Swedish Research Council through grant agreements no. 2022-06725 and no. 2018-05973. Finally, we thank the sheep research community and specifically the authors of our sources for making their genotype and sequence data available to the public.

## DATA AVAILABILITY

New raw sequence data for the ancient individuals are available through the European Nucleotide Archive (ENA) under project accession number PRJEB105076. Genotype calls can be found on Zenodo (10.5281/zenodo.18470345).

## Materials and Methods

### Ancient DNA extraction and Sequencing

Tooth and jaw bones were recovered from 16 ancient caprine individuals from the El Portalón de Cueva Mayor and 8 from Els Trocs, two caves in Northern Iberia (for details on each site, see Supplementary Note 1). DNA extraction and library preparation were done at the dedicated aDNA facilities of the SciLifeLab Ancient DNA Unit at the Department of Organismal Biology, Uppsala University, Sweden. Before extraction, samples were irradiated with UV light (6 J/cm^2^ at 254 nm) for 20 minutes, and then their outer surface was removed using a Dremel drill. Samples were then wiped with sterile cotton swabs with 0.5% sodium hypochlorite solution and UV-irradiated Mili-Q water, and exposed to UV-irradiation again on each surface. Then sub-samples of 50-100 mg were cut from each sample using a Dremel tool. DNA was extracted using a modified silica-based method (1). Instead of sodium dodecyl sulphate, 1 M urea was used in the extraction buffer. Every 10 samples, one extraction blank was added as a negative control. Subsamples were pre-treated with 1 ml of 0.5 M EDTA (Invitrogen) and incubated at 37°C for 30 min. The EDTA solution was removed and the subsample was digested with 1 ml extraction buffer (0.44 M EDTA/1 M urea) containing 0.25 mg/ml proteinkinase K (Sigma-Aldritch). They were incubated with rotation for ∼ 23 h at 37°C and then at 55°C for 6.5 h. The supernatant was collected and stored at − 20°C. 1 ml of fresh extraction buffer with proteinkinase was added to the sample and incubated further at 55°C for 19 hours. Supernatants were combined and concentrated using an Amicon Ultra-4 filter unit (Millipore). DNA was purified using MinElute PCR purification kits (Qiagen) and eluted in a total volume of 110µl EB buffer. Qubit dsDNA HS assay (Invitrogen) was used to determine the concentration of the DNA extract.

Doubled-stranded blunt-end libraries were prepared from 20µl of DNA extraction according to protocol, with some modifications (2)(MineElute PCR purification kits were used to clean enzymatic reactions instead of SPRI beads). For every 10 samples, an extraction and library blank was added as a negative control. Libraries were quantified by real-time qPCR in a 20µl reaction using Maxima SYBR green master mix (Thermo Fisher Scientific), 200 nM IS7 primer and 200 nM IS8 primer, to determine the number of indexing cycles. Dual-indexing PCR amplification was performed in duplicates in a 50µl reaction using 6µl DNA library, 5 U AmpliTaq Gold DNA polymerase (Thermo Fisher Scientific), 1 × GeneAmpl Gold Buffer (Thermo Fisher Scientific), 2.5 mM MgCl_2_, 250µM of each dNTP, 200nM P7 indexing primer and 200nM P5 indexing primer (2, 3). PCR reaction was performed at 94°C for 10 min, 13-20 cycles of (94°C for 30 s, 60°C for 30 s, 72°C for 45 s) and 72°C for 10 min. Duplicates were pooled and purified with AMPure XP beads (Beckman Coulter). The quality of the libraries was analyzed by the 2200 Tapestation System (Agilent) and the Qubit dsDNA HS assay (Invitrogen) was used for quantification of the sequencing libraries.

For individuals where both jawbone and teeth were available, two extractions were performed. The libraries were initially sequenced on an Illumina NovaSeq 6000 SP lane using 150bp paired-end reads at the SciLifeLab SNP&SEQ Technology Platform at Uppsala University. Of the 16 El Portalón samples, one was discarded for poor preservation of the bone before extraction. A first round of 10 samples from El Portalón was screened for sheep DNA, and a second round of screening was performed with 5 samples from El Portalón and 8 from Els Trocs. All samples belonged to sheep with the exception of a Roman goat from El Portalón, which has been included in another study (4). Eight sheep individuals from El Portalón with over 1% of endogenous DNA were subjected to deeper resequencing. Resequencing was performed using an Illumina NovaSeq 6000 S4 flow cell with 100bp paired-end reads and v1 chemistry at the SNP&SEQ Technology Platform in Uppsala.

### Next Generation Sequencing Data Processing and Authentication

In order to verify that the sequences belonged to sheep, we used *FastQ Screen* (5), which maps the input sequences to a set of reference organisms. All samples included in downstream analysis showed a significantly higher proportion of sequences identified as sheep than any other species in the panel, including goat, mouse, rat, and human.

Paired-end reads were then merged using *FLASH 1.2.11*(6) and the remaining adapters were removed using *CutAdapt 2.3* (7) set for NovaSeq sequences (*–nextseq-trim* 15) and a maximum error rate of 0.2. Trimmed and merged sequences were then mapped to the Texel sheep reference genome 4.0 (Oar4.0) with the mitochondrial sequence from NCBI accession number NC_001941.1 using *BWA aln 0.7.17-r1188* (8) with parameters *-n* 0.01 *-o* 2 and seeding disabled. Aligned sequences with mapping quality below 30 were then filtered using *SAMtools view* and then sorted *SAMtools sort* (9). Potential PCR duplicates were removed using *DeDup 0.12.8* (10). Overall mapping quality was assessed using *Qualimap 2.2.1* (11), and damage patterns consistent with aDNA deamination were observed with *MapDamage 2.0.8* (12) (Supplementary Figure 1). Then, we used *ANGSD 0.937-92-g5cd355b* (13) to create pseudo-haploid calls by picking one of the bases at random for each position from a set of previously defined sites ascertained in wild sheep (see below) using the *-sites* command and with settings *-dohaplocall 1 - doCounts 1 -doGeno -4 -doPost 2 -doPlink 2 -minMapQ 30 - minQ 30 -doMajorMinor 1 -GL 1 -domaf 1*. Transitions were removed during this step using the *-rmTrans 1* setting.

Due to the low amount of data, samples from El Portalón and Els Trocs with less than 0.01× coverage were excluded from (most) downstream analysis. This left a total of 11 samples from El Portalón from the Neolithic to the Roman period and 6 samples from Els Trocs, belonging to the Early to Late Neolithic.

### Modern Corsican mouflon DNA extraction, sequencing and data processing

One Corsican mouflon female’s genome was also sequenced for comparison. DNA was extracted from whole blood using the GenElute™ Mammalian Genomic DNA Miniprep Kit following the standard protocol. A sequencing library was prepared using a ThruPlex DNA-seq kit and sequenced as one out of 26 libraries on an Illumina NovaSeq 6000 S4 lane with 150bp paired-end reads at the SciLifeLab SNP&SEQ Technology Platform at Uppsala University. The sequence data showed indications of fragmentation, with most fragments shorter than 300bp. Therefore, the paired-end reads were then merged with *FLASH 1.2.11* (6) and adapters were removed using *CutAdapt 2.3* (7) set for NovaSeq sequences (*–nextseq-trim* 15), and both the combined and not combined reads were mapped using *BWA-mem 0.7.17-r1188* (8) to the same reference as the ancients. Mapped reads were then mate-scored using *samtools fixmate* (9) and recoded into bam format with *samtools view* and settings *-bhu -F 4*. Once in bam format, reads were sorted with *samtools sort* and duplicates were removed using *samtools markdup -r*. Then, bam files were merged into one with *samtools merge*. Variants were called using *bcftool 1.14 mpileup* (14) with standard settings and variants were then filtered *Qual >*= 20 and *MQ >*= 30, and restricted to the sites listed in the reference SNP panel (see below).

### Molecular sex determination and mitochondrial haplotyping

As the mapping assembly does not contain the Y chromosome, biological sex was determined by comparing the average read depth of the autosomal and the X chromosomes using *SAMtools idxstats* and used the R_x_ ratio (15). Samples with similar coverage for all chromosomes were assigned as females, while those with half the amount of sequences mapping to the X chromosome were assigned as males (Supplementary Table 1). One sample (APOR-001) could not be assigned a biological sex due to a very low proportion of endogenous DNA.

We used the *Mapping Iterative Assembler* (MIA) *5a7fb5a* to call consensus sequences of the mitochondria of the ancient samples. To avoid reference bias, we used the Asiatic Mouflon mitochondrial sequence (NCBI accession NC_026063), along with a specific substitution matrix suited for aDNA damage. Consensus sequences were assembled for sites with coverage higher than 3x and 10x as cutoff values using Mapping Assembler in MIA. Sequences were aligned using *MAFFT 7.407* (16) with standard settings and visually inspected. Haplotypes were called from those sequences using *MitoToolsPy-Seq* from *MitoTool 943ce25* (17) using the sheep profile and the whole mitochondrial sequence.

### Radiocarbon dating

Samples for which there were sufficient amounts of remaining material after sampling for DNA were radiocarbon-dated by the Tandem Laboratory at Uppsala University or the National Laboratory of Age Determination at the NTNU University Museum in Trondheim. Bones and teeth at Uppsala were mechanically cleaned by scraping the surface and then ground in a mortar. 0.25*M* HCl was used for incubation at ambient temperature for 48 hours. 0.01*M* HCl was added to the insoluble fraction and incubated at 50°*C* for 16 hours. The soluble fraction was added to a 30*kDa* ultrafilter and centrifuged, and the retentate was lyophilized. Before determination, the fraction to be dated was combusted to CO_2_ using a Fe-catalyst. For the material radiocarbon-dated at Trondheim, the protocol was as described in Seiler et al, 2019 (18). All acquired dates were calibrated with *OxCal 4.4* (19) using *IntCal20* (20) as the calibration curve.

### Modern reference data

To avoid possible bias induced by modern selection and breeding practices, we used a modern reference panel of 188 published modern sheep genomes, where only Single Nucleotide Polymorphisms (SNPs) ascertained in nine individuals from six different wild sheep species and a goat collected from dbSNP (Supplementary Table 3) (21–24) were used as in Katpan et al, 2024 (25) and Larsson et al. 2023 (26). Genomes for all samples were mapped with *BWA mem* (8) and then converted to bam format and sorted using *SAMtools* (9) *view* and *sort*. Duplicates were removed with *Picard MarkDuplicates* (27) and reads with mapping quality lower than 20 were filtered using *SAMtools view*.

Once mapped, GVCF files for the wild reference individuals were created with *GATK HaplotypeCaller* (filtering for *MQ >*= 30) (28) for each sample. ThenSeven Thousand Years of interwoven Genomic History between Sheep and Humans in Iberia they were combined with *GATK CombineGVCFs* and genotypes were called using *GATK GenotypeGVCFs*. Called SNPs were filtered using *bcftools view* (14, 29), keeping only bialellic snps with *Qual >*= 30, *QD >*= 2, *SOR <*= 3, *FS <*= 60 and *MQ >*= 40. Files were recoded to VCF format using *vcftools* (14, 29), also filtering sites with minor allele frequency (MAF) of 0.05 and excluding sites not in Hardy Weinberg Equilibrium (HWE) with a threshold of 0.001. SNPs in the domestic dataset were called using GATK *HaplotypeCaller* on the positions ascertained on the wild SNP file and then filtered with *bcftools view* (14) to allow only biallelic SNPs with *Qual >*= 30, *QD >*= 2, *SOR <*= 3, *FS <*= 60, *MQ >*= 40, *M QRankSum >* − 12.5 and *ReadP osRankSum >* 8. The GVCF file was then recoded using *vcftools*, allowing only sites in HWE with a threshold of 0.001. Then both files were converted into *Plink* bed format using *Plink2* (30), filtering out any SNP with missingness higher than 0.05 and any sample with higher missingness than 0.1. Then both datasets were merged using the *–bmerge-files* on *Plink 1.9* (31). The resulting dataset comprised 200 samples and 4, 603, 349 SNPs.

To merge ancient and modern data, files were converted back to VCF format with *Plink 1.9 –recode vcf* and merged using *bcftools merge* (14, 29) with standard settings and then converted back to bed and tped format using *Plink 1.9*. To avoid technical issues with some methods, a dataset with all sites pseudo-haploidized for all samples was created by randomly picking one of the calls.

### Exploratory population genetic analysis

This haploid dataset was used to explore the population structure of ancient Iberians. Principal Component Analysis (PCA) was performed using SmartPCA from the EigenSoft package (32, 33) with standard settings and lsqproject activated for all the ancient samples and after pruning our dataset using *Plink 1.9* with the parameter *–indep-pairwise 50 5 0.2* (31). This pruned data was also used to run an ADMIXTURE (34) analysis using version 1.3.0, where the number of ancestry components (*K*) ranged from 2 to 15 and each *K* was run 20 times. ADMIXTURE results were then compared, plotted and visually inspected using Pong (35).

### Describing ancient Iberian sheep demographic history

As we wanted to describe the demography of ancient Iberian sheep, we ran OrientAGraph (36) on a sub-sample of the haploid dataset that only included the highest quality ancient samples per period (Supplementary Table 3), where only SNPs with no missingness were kept. As proxies for European and Asian populations, samples from the Altay and Merino breeds were selected. This led to a dataset of only 126491 SNPs and 28 samples. OrientAGraph was run with the Asiatic Mouflon as root, a seed of 12345 and the *-allmigs -mlno* and *-noss* parameters and allowing for 0 to 6 migration edges. Admixture graphs were then plotted in *R 4.2.2* (37) using *Treemix* (38) plotting functions.

We also used qpGraph (39) as implemented in *Admixtools2* (40) to reconstruct admixture graphs with the find_graphs() function. We first generated an *f*_2_ matrix from the genotype data with the settings maxmiss=0.15 and auto_only=FALSE. Next, we used find_graphs() to systematically explore possible admixture graphs with stop_gen = 10000, stop_gen2 = 30 and plusminus_generations = 10 as suggested in (41). For between 0 and 6 migration events, we started the graph search with 500 different random seeds and the constraints that modern Asian sheep (represented by Altay), El Portalón Neolithic, European Mouflon and Anatolia Neolithic (represented by tps062) were unadmixed. Graph fits for different numbers of migrations were then compared using qp-graph_resample_multi() and compare_fits() employing 1000 bootstraps.

To assess the relationship of ancient Iberian sheep with other ancient and modern domestic sheep we estimated Outgroup-*f*_3_ statistics (39) using *Admixtools2* (40) in following the format *f*_3_(*Asiaticmouflon*; *non* − *Iberiansheep, ElP ortal*ó*n*), where the El Portalón samples were divided by period and compared against each other in the form of biplots. We used *f*_4_ statistics to assess the relationship between the different periods from El Portalón with Eastern Eurasian breeds using the configuration of *f*_4_(*Asiatic Mouflon, Altay*; *ElP ortal*ó*n, ElP ortal*ó*nN eolithic*), using APOR-008 (our oldest sample from that site) as a Neolithic baseline where “El Portalón” refers to all other samples from that site. This analysis was mirrored on the human dataset from the Allen Ancient DNA Resource (AADR) V54.1.p1 (42) such that *f*_4_(*Chimp, Y amnaya, Iberian, IberianN eolithic*), using the oldest Iberian Neolithic human in the dataset (CB13) (Supplementary Table 4). We also calculated *f*_4_ statistics to test if there was a recent influx of Eastern ancestry in European breeds with *f*_4_(*Asiatic Mouflon, Altay*; *M odernBreeds, ElP ortal*ó*nRoman*). Changes in genetic diversity over time were measured by estimating the expected heterozygosity at pre-ascertained transversion sites *popstats –pi*(43). This function calculates the mismatch probability of a given SNP site between two random samples from a given population. For El Portalón, samples were divided by temporal range (e.g. Neolithic, Pre-4.8k Chalcolithic, Post-4.8k Chalcolithic and Bronze Age). For APOR-013, the Roman period sample, we used an ultra-low coverage Roman sample, APOR-001, which allowed for the comparison at 2531 SNP sites. Other ancient and modern populations were analyzed as a whole.

## Supplementary Note 1: Sites and Samples

### El Portalón de Cueva Mayor

This rich archaeological site is located in the karstic system of the Sierra de Atapuerca complex in Burgos, Northern Iberia. Archaeological excavations at this site have unearthed a stratigraphic sequence that extends from the Late Pleistocene to the Holocene up to the Early Middle Ages. At this site, more than 100 direct radiocarbon dates from different strata range from 30000 to 1000 cal BP (44, 45). Although the Late Pleistocene strata display scarce archaeological activity, the Holocene period experienced intense human activity almost uninterruptedly from the Mesolithic to Medieval times. Abundant archaeological artifacts belonging to all chronocultural periods have been found. Although human remains have been found throughout the chronostratigraphy, domestic faunal remains are more common, suggesting that over most of its use, this cave housed livestock, with short periods of human occupation. Early Neolithic fauna recently recovered is characterized by hunted taxa, which changes in the Late Neolithic to a predominance of domestic taxa. Caprines, followed by cattle and pigs, are the most abundant domestic species from the Neolithic to the early Bronze Age. This pattern changed by the Middle Bronze Age, when cattle became more abundant than caprines (46–48). Archaeological evidence suggests these animals were exploited as livestock (48), but some remains have also been found as part of cultural activities (e.g. grave goods associated with a Chalcolithic child burial, described in (45)).

### Els Trocs

Els Trocs cave is located on the southern slope of a conical hill at an altitude of 1500 meters at the village of San Feliu de Veri/Bisaurri, in the Pyrenees of Huesca. The proximity to high-altitude pastures and to saline springs has made this area a strategic location for transhumance in historic and probably throughout prehistoric times. 2000 years of human occupation of this site have been described based on its stratigraphic sequence and direct radiocarbon dates, with six different phases of occupation (49):

- Trocs Ia. The first phase of occupation of the cave, dated to the last third of the 6^th^ millennium cal. BC. This layer is characterized by a ceramic vessel floor and the presence of 9 human individuals buried on top. These individuals were closely related and were killed in a display of extreme violence (50).
- Trocs Ib. The partial caving of the entrance 200 years later meant the end of the ritual phase of the early habitation of Els Trocs. The humans occupying the cave during this period dug 2 trenches in the ground and used the displaced earth to cover the remains that had previously been displayed for ritual purposes. Remains from this period belong to fauna, especially caprines. Most of the sheep samples that come from this site presented in this study (ATRO-004 to ATRO-008) belong to this phase.
- Trocs II. Dated to the middle of the 5^th^ millennium cal. BC. Characterized by the presence of amortized hearths and the seasonal use of the cave by transhumant shepherds. ATRO-003 belongs to this phase.
- Trocs III. From the first third of the 4^th^ millennium to the early beginnings of the 3^rd^ millennium cal. BC, characterized by smaller habitations, is believed to be designed for short occupations. There is also an area of the cave at this period that is used for funerary purposes.
- Trocs IV. Partial funerary deposit from the transition between Neolithic and Chalcolithic (3^rd^ millennium cal. BC). ATRO-001 and ATRO-002 belong to this phase.
- Trocs V. Last occupation of the site on the most superficial phase of the stratigraphy. Imperial Roman remains have been found (4^th^ century BC).

Dating of the different occupations is supported by a series of radiocarbon dates carried out for each phase of the site (49, 51, 52).

## Supplementary Note 2: Population genomic analysis

### Population affinities

Population affinity changes between different time periods at El Portalón were compared through biplots of their respective Outgroup-*f*_3_ statistics (39) with the same set of modern and ancient groups (Figure 2, Supplementary Figure 3). The biplot in Figure 2 summarizes all periods compared to the Post-4.8k Chalcolithic samples, while Supplementary Figure 3 represents half of a matrix of pair-wise comparisons between ages.

Our results show higher affinity of the Neolithic Iberian sheep with the European mouflon population in all comparisons, while later periods show significantly higher affinities with modern domestic breeds. The affinities to the different ancient populations, however, differ depending on the comparison. In the comparison between Iberian Neolithic and Pre-4.8k Chalcolithic sheep, both Iranian and Anatolian Neolithic are not significantly different between the time periods, as their confidence intervals overlap with the *x* = *y* diagonal. The Caucasus Bronze Age sample, while displaying a higher affinity to the Pre-4.8k Chalcolithic, has wider error bars, making this comparison also not significant. The only ancient sample that is barely significant is the Scandinavian sheep from Stora Forvar (ASTF-001). These results are quite similar to the Bronze Age versus Neolithic comparison, with all ancient populations sharing similar amounts of drift with both periods, with the exception of the Anatolian Neolithic, which have more affinity with the Neolithic sheep from El Portalón.

This trend, however, seems to have been reversed during the Post-4.8k Chalcolithic, when we see a substantial increase in affinity of the Caucasus Bronze Age sample, and more modest increases in the affinity of the Scandinavian Neolithic sheep, which previous studies already suggested was admixed with some Eastern/Central Asian ancestry (25, 53). This higher affinity with the Caucasus sample can be seen in all other comparisons against this time period.

The comparison between the Bronze Age with other periods displays similar patterns to those of the Pre-4.8k Chalcolithic, and the comparison between these two only shows small differences in the Neolithic sheep from Iran and Anatolia, and the European Mouflon. The comparison between the Bronze Age sheep and the Post-4.8k Chalcolithic, however, is quite interesting, as it is the only time modern domestic sheep show significantly higher affinities to the older of the two time periods compared. These suggest the Bronze Age population displayed a more “ancestral” gene pool (e.g. a population related to the Pre-4.8k Chalcolithic that was not admixed with the Post-4.8k Chalcolithic one) or that modern breeds, including the Iberian sheep, are more related to the population that was the source of the increased Caucasus affinity.

Lastly, the Roman individual displays a significantly higher affinity with modern sheep across most comparisons. When compared with the Neolithic and Pre-4.8k Chalcolithic, only the European mouflon shows a higher affinity with the older sheep, which supports their classification as feral Neolithic sheep. Interestingly, the Anatolian Neolithic samples are not significantly different, while all other populations show higher affinity to the Roman population. This pattern, however, is broken during the Post-4.8k Chalcolithic. There, we see that the Post-4.8k Chalcolithic sheep still have higher affinities with the Caucasus Bronze Age, in addition to the European mouflons and Anatolian Neolithic sheep. The Roman sample, on the other hand, has higher affinities with Asian, Middle Eastern and African modern domestics, and most of the European production breeds. Archaic European breeds and the Iranian and Scandinavian Neolithic samples are not significantly different.

### Admixture graph reconstruction

#### qpGraph

We explored admixture graphs (39–41) with up to six admixture/migration events (Supplementary Figure 5). In addition to the Iberian groups, we included European mouflon, Anatolian Neolithic (TPS), a representative of modern Asian sheep breeds (Altay), Bronze Age Russians (ACau), Late Neolithic Scandinavia (ASTF), and Asiatic mouflon as an outgroup. We are discussing the best-fitting model for each complexity level below.

Without allowing for any admixture (Supplementary Figure 5a), the structure first follows geography (with an eastern and a western main branch). In the western branch, Neolithic Anatolia first splits off, followed by ASTF, Roman Iberia and Bronze Age Iberia. The European mouflon groups with Iberian Neolithic in a sister clade to the two Chalcolithic groups.

The first migration event models ASTF as a mix of Eastern and Western ancestries, the Eastern ancestry originates in a population related to Altay, the Western ancestry is related to Roman Iberians (Supplementary Figure 5b).

The model with two migration events groups Roman Iberians and ASTF, as an admixed group with basal Western ancestry and Eastern ancestry related to ACau (Supplementary Figure 5c). The second admixture event models Chalcolithic and Bronze Age Iberians as a mix between Roman Iberians and Neolithic Iberians.

Adding a third migration event results in a model where Roman Iberians receive an additional pulse of ancestry, modelling them as admixed between Bronze Age Iberians and a population distantly related to ASTF (Supplementary Figure 5d). Furthermore, the Eastern ancestry in ASTF and Chalcolithic Iberians comes from the same source, which is related to ACau.

The fourth migration event adds a pulse of western ancestry to ACau (Supplementary Figure 5e). More complex models account for slight differences in Iberian groups with Bronze Age Iberians receiving additional Eastern ancestry (m=5; Supplementary Figure 5f) and Pre-4.8k Chalcolithic Iberians receiving additional basal western ancestry relative to Post-4.8k Chalcolithic Iberians (m=5; Supplementary Figure 5g). Comparing the model fit across different complexities (Supplementary Table 6) suggests that one migration event is a significantly better representation of the data than a tree model without admixture (*p* = 0.0487), but two migration events did not provide a better fit than one (*p* = 0.179). We still decided to display a model with three migration events in the main text due to several reasons:

- The best-fitting models for m=1 and m=2 are using the Roman Iberian sample as a source for Eastern ancestry in other groups, which violates the temporal order of the samples, as this is the most recent of all ancient samples in this study.
- At m=1, Eastern ancestry in Scandinavia is modeled, which is not directly relevant to Iberia. m=2 adds Eastern ancestry in the Roman Iberian group but not in Prehistoric groups. As our other results point to at least one additional gene flow event into Iberia during the Chalcolithic, we believe that a third migration event is justified in this model.
- The model with three events is a significantly better fit to the data than a model with one event (*p* = 0.00773). More complex models are either not significantly better fits than m=3 (as for m=4; *p* = 0.132) or are only marginally better (m=5; *p* = 0.047 and m=6; *p* = 0.046)

Overall, one can assume that the qpGraph model with three migration events is still an oversimplification, as it summarizes all prehistoric Eastern ancestry into a single event. Our other results, however, suggest that there were two or three separate events during the Chalcolithic and Bronze Age, some of which are minor and difficult to differentiate in a graph model, considering the availability of reference data and their genome coverage.

#### OrientAGraph

OrientAGraph (36, 38) is another method for constructing admixture graphs. We again included the Iberian temporal groups, European and Asiatic mouflon, as well as Altay as a proxy for Asian sheep and Merino as a proxy for modern European sheep breeds. Due to concerns about missing data, the Anatolian Neolithic, the Russian Bronze Age, and the Scandinavian Late Neolithic samples were not included in this analysis.

The tree structure without any gene flow events is similar to the qpGraph model with the same complexity (Supplementary Figure 5A). The first migration edge is between European and Asiatic mouflon, not changing the topology otherwise or affecting any of the ancient groups (Supplementary Figure 5B). The second migration edge adds gene flow from the domestic European breeds to the European mouflon (Supplementary Figure 5C), which is in line with previous results (54). The third migration edge establishes a connection between the Roman sample and Asian breeds (Supplementary Figure 5D), similar to the signal seen in qpGraph and other analyses that the Roman Iberian sample contained most Eastern ancestry.

The fourth migration edge adds basal ancestry to the common ancestor of Iberian Neolithic and Pre-4.8k Chalcolithic as well as the European mouflon (Supplementary Figure 5E). Consequently, the Bronze Age and Roman groups do not receive this pulse, meaning they are modeled as samples with less basal ancestry but a gene pool more similar to the modern breeds. The fifth migration edge connects the Neolithic Iberians with modern European breeds (Supplementary Figure 5F) while the sixth migration edge adds a minor pulse of additional basal ancestry into the Bronze Age Iberians (Supplementary Figure 5G), again suggesting a slightly different ancestry of the Bronze Age samples compared to older time periods.

qpGraph and OrientAGraph are not completely consistent in their modeled migration events. This can be partly explained by their different approaches of handling missing data and, consequently, the different groups used in the two analyses. Nevertheless, both of them consistently suggest multiple pulses of additional Eastern ancestry into Iberian sheep.

## Supplementary Figures

**Supplementary Figure 1.**
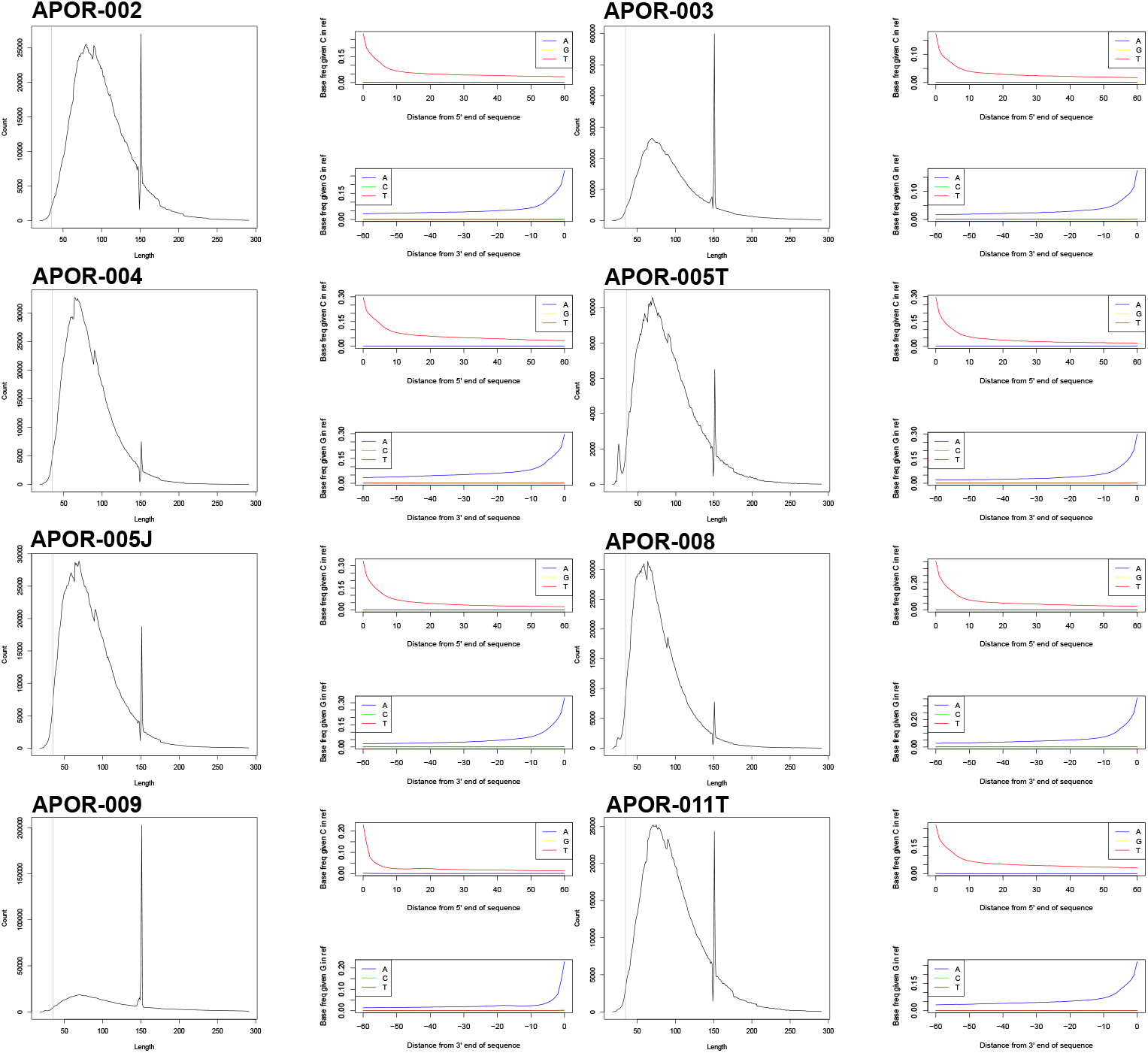
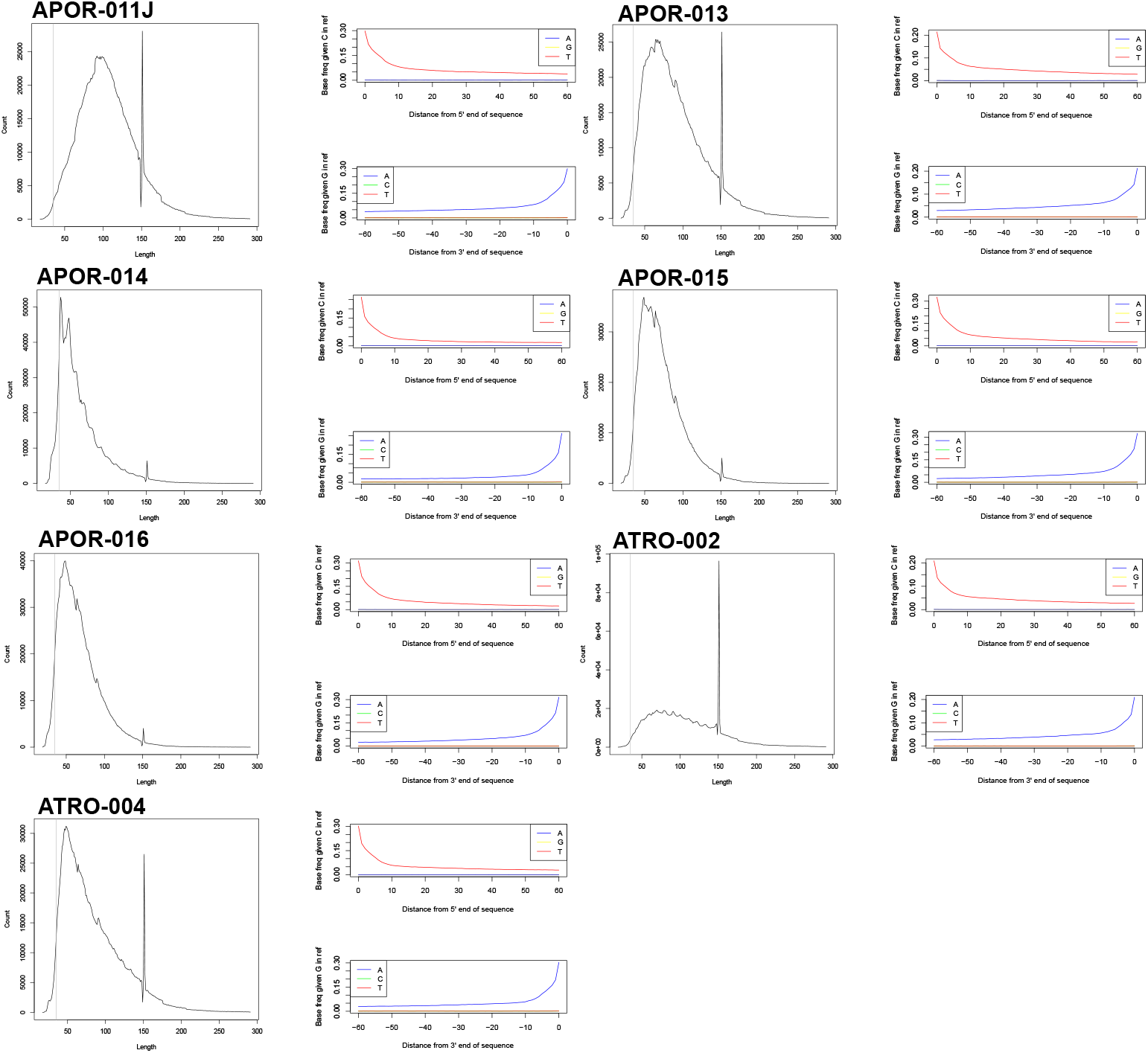
Fragment lengths and deamination plots per sample.

**Supplementary Figure 2.**
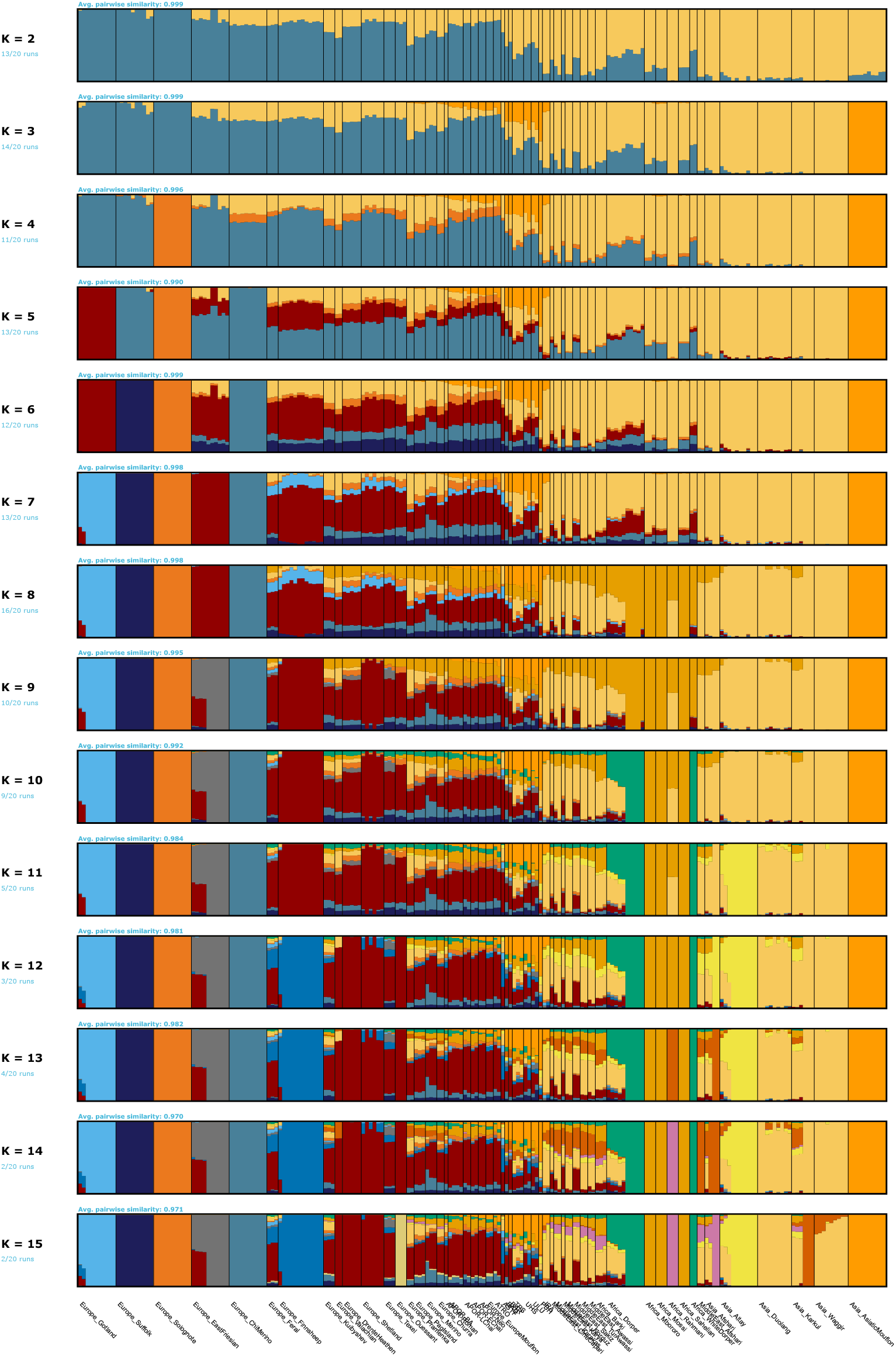
ADMIXTURE results for K=2-15

**Supplementary Figure 3.**
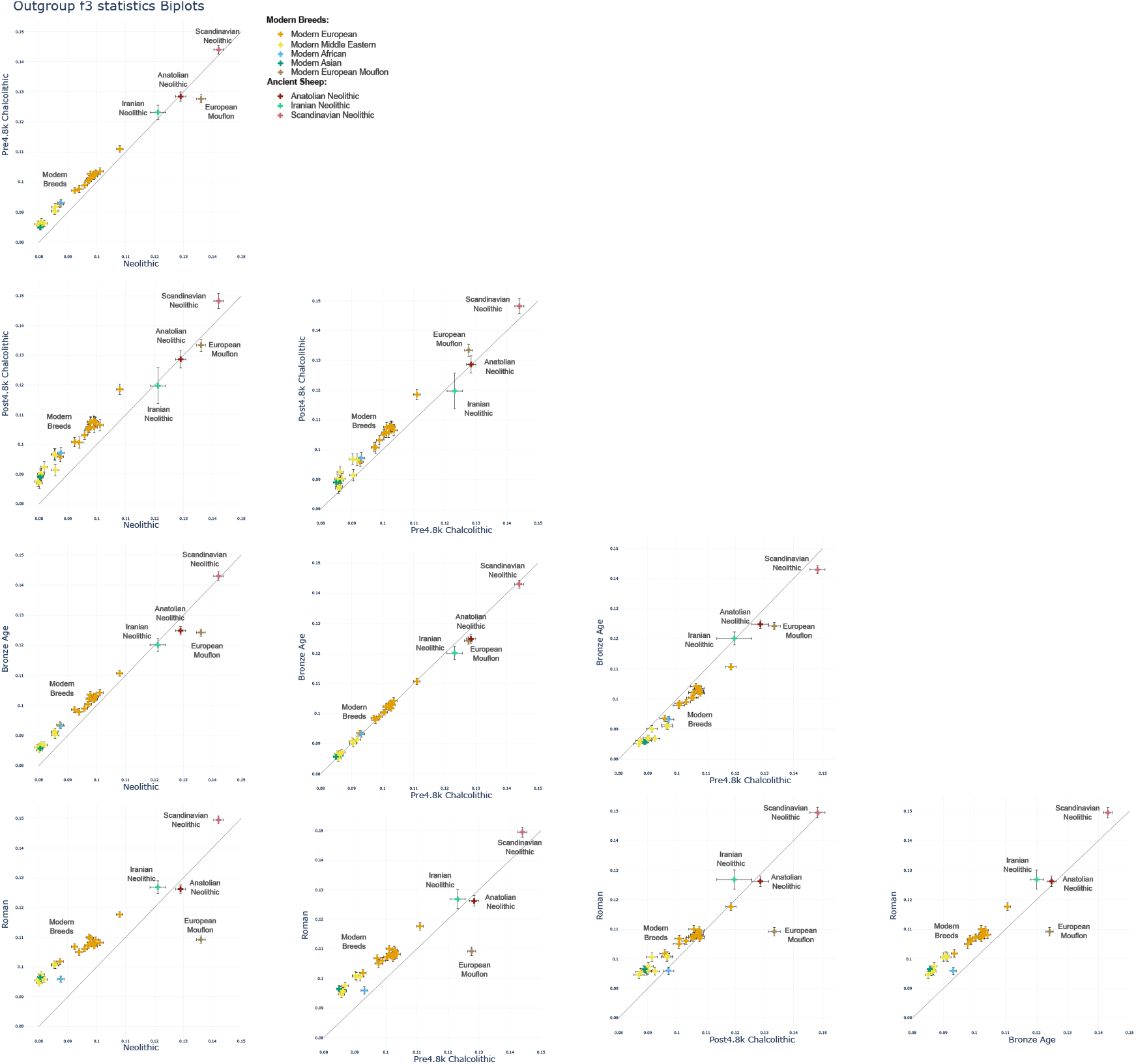
Biplot matrix comparing the Outgroup-*f*_*3*_ statistics of every combination of periods.

**Supplementary Figure 4.**
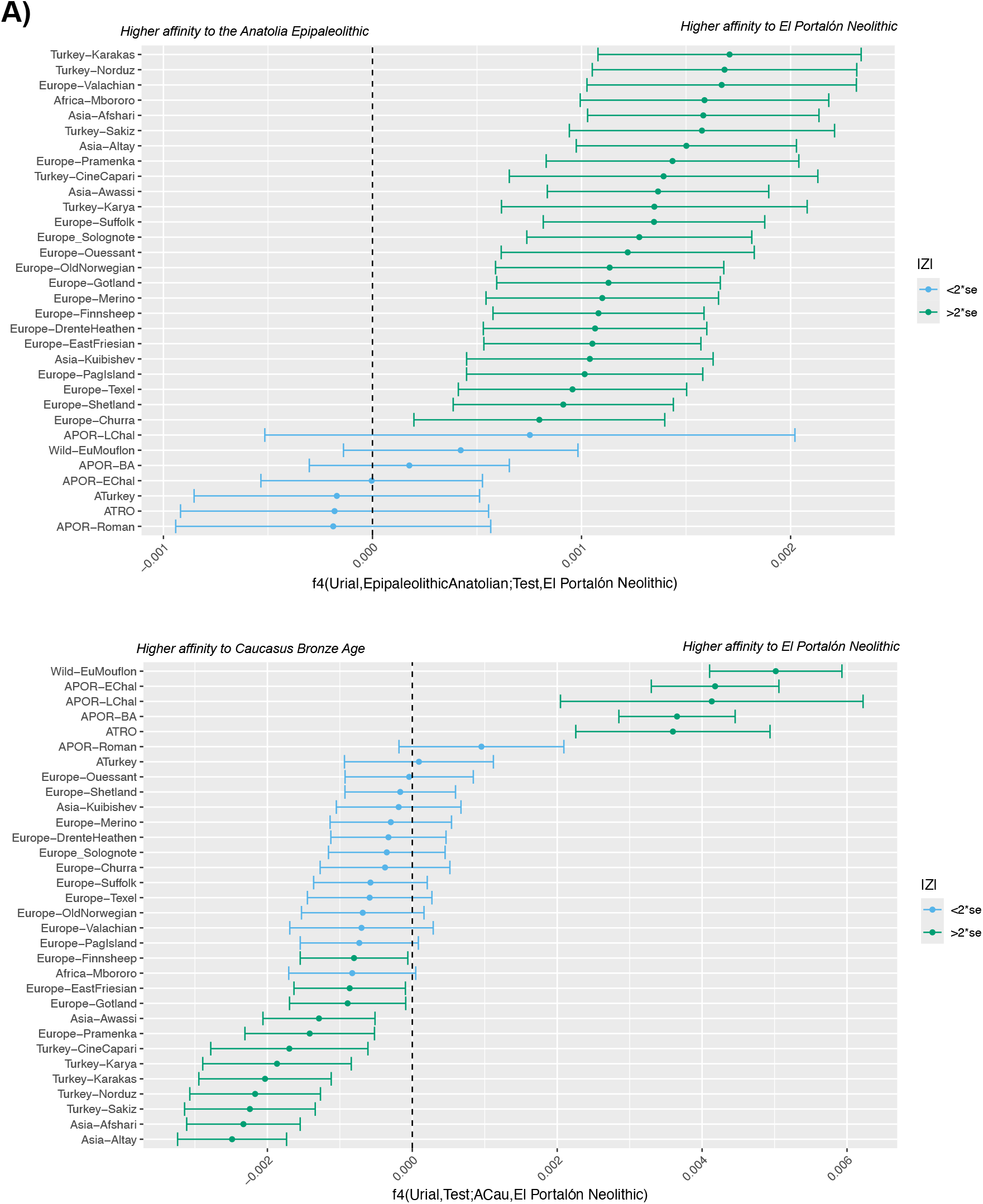
A) F4 statistics in the shape of f4(Urial,EpipaleolithicAnatolian;test,El Portalón Neolithic). B)F4 statistics in the shape of f4(Urial,test;BronzeAgeCaucasus,El Portalón Neolithic). A) shows the European Mouflon as the only modern population that is equidistant between the Epipaleolithic Anatolian and Iberian Neolithic samples. B) shows that the European Mouflon and the prehistoric Iberian samples have a stronger affinity to the El Portalón Neolithic than the sample from the Caucasus, while modern sheep from Asia show significant affinity with the Caucasus sample. APOR-EChal and APOR-LChal refer to the Pre-4.8k and Post-4.8k Chalcolithic samples, respectively.

**Supplementary Figure 5.**
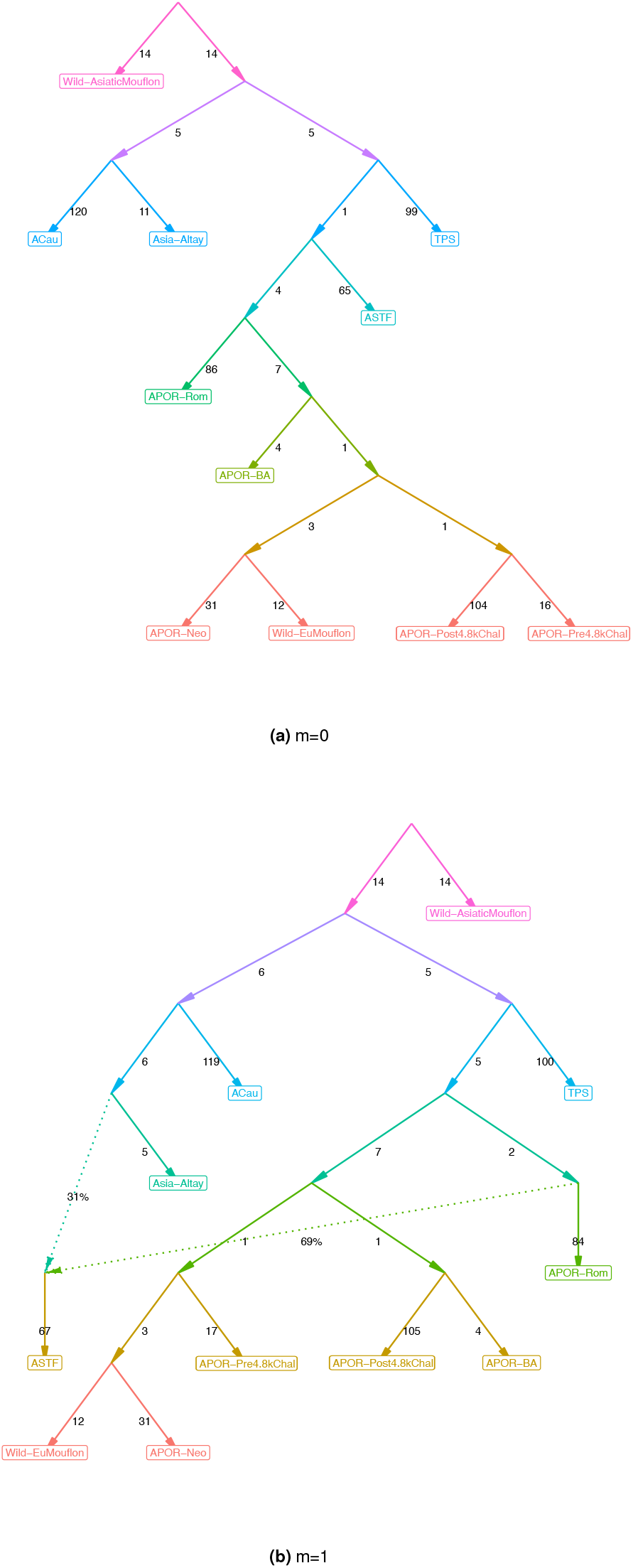
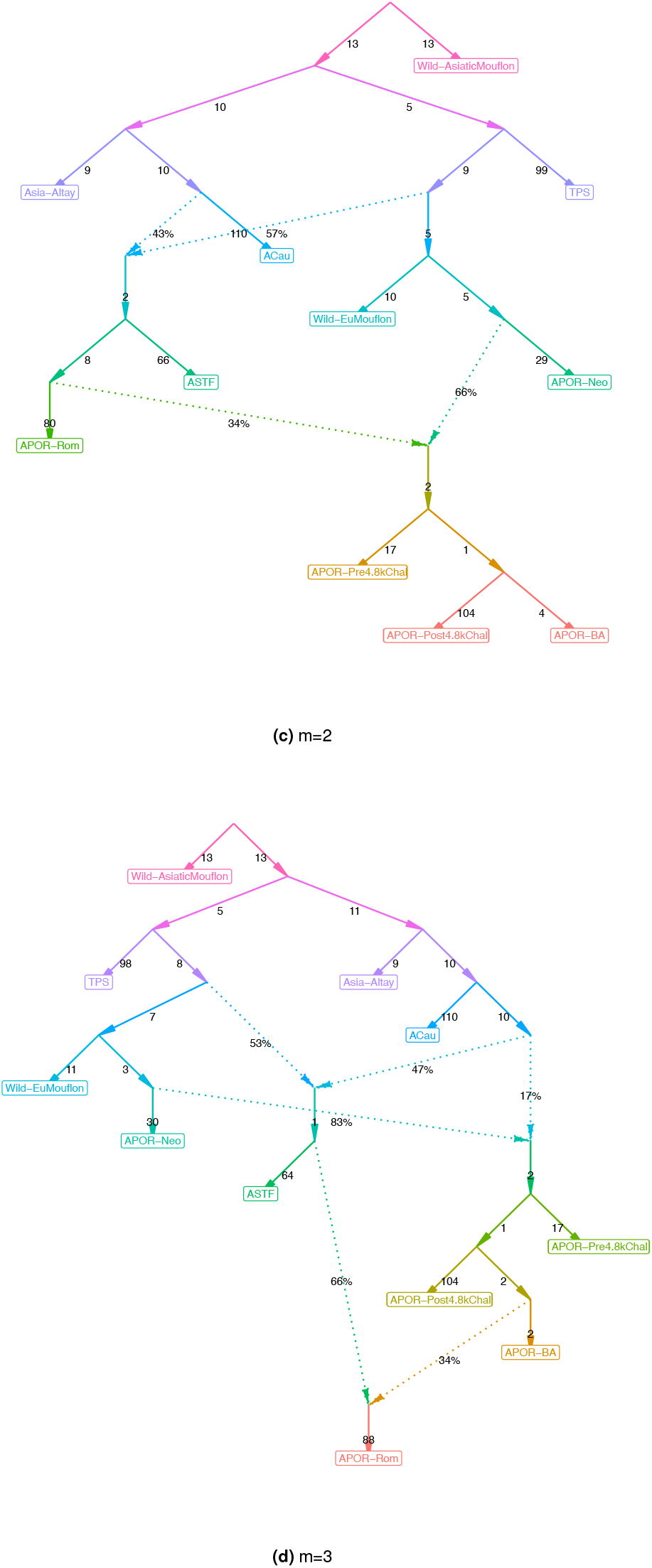
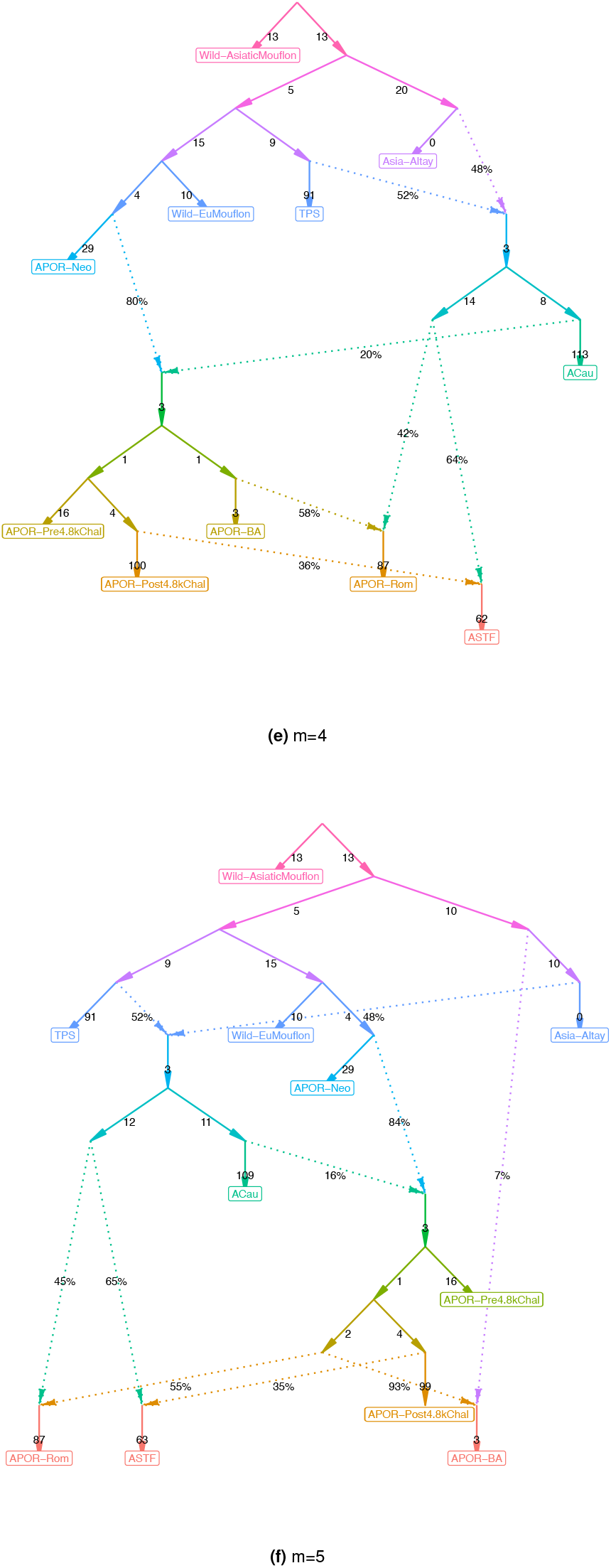
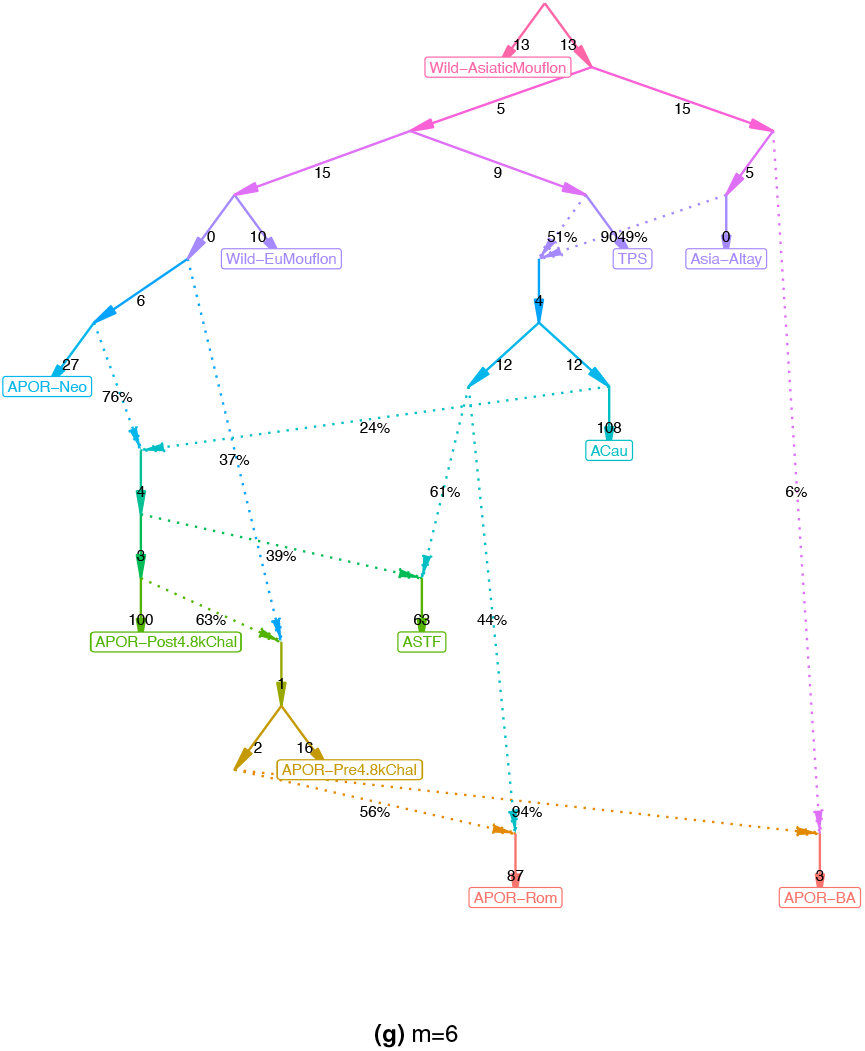
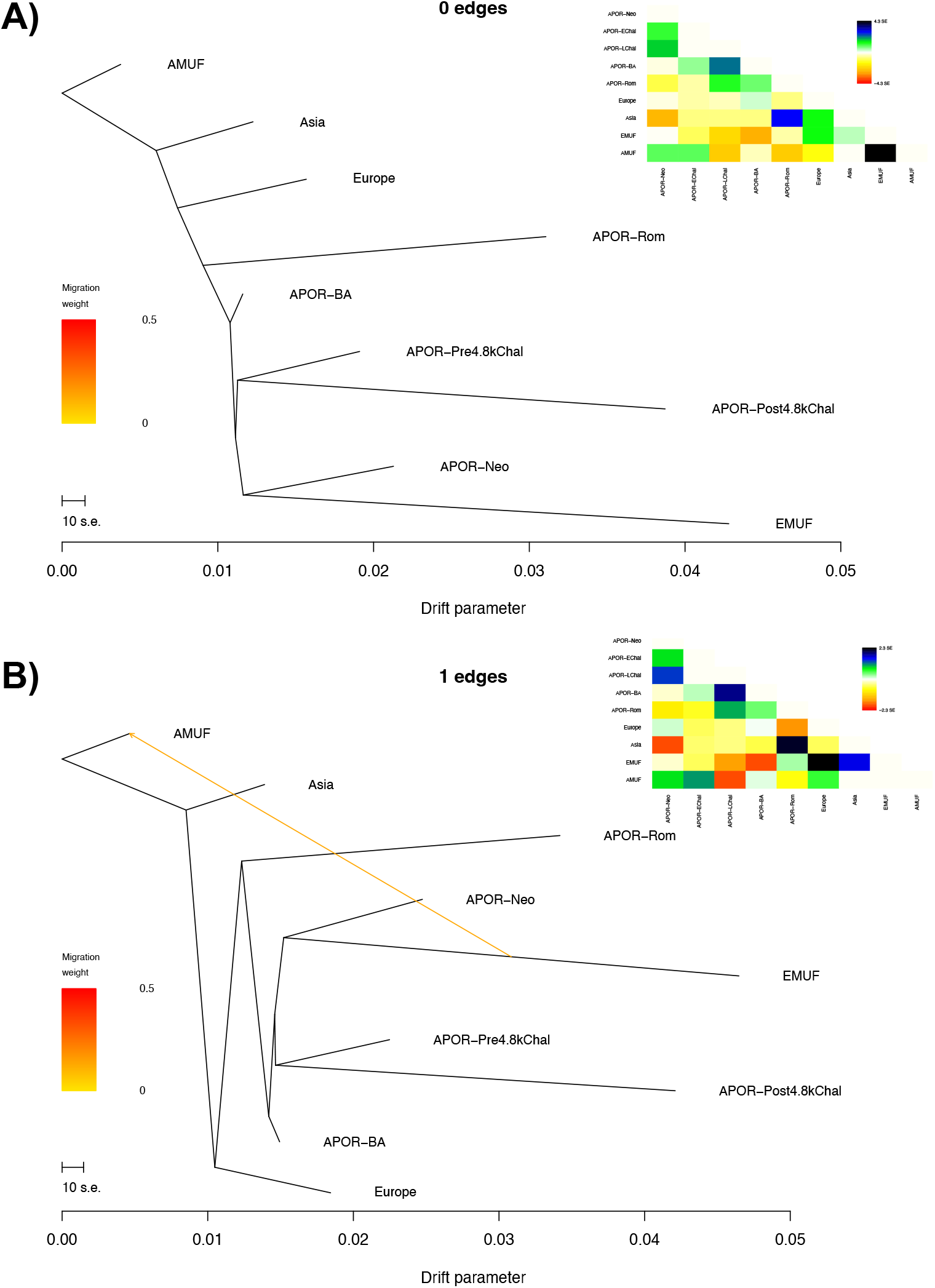
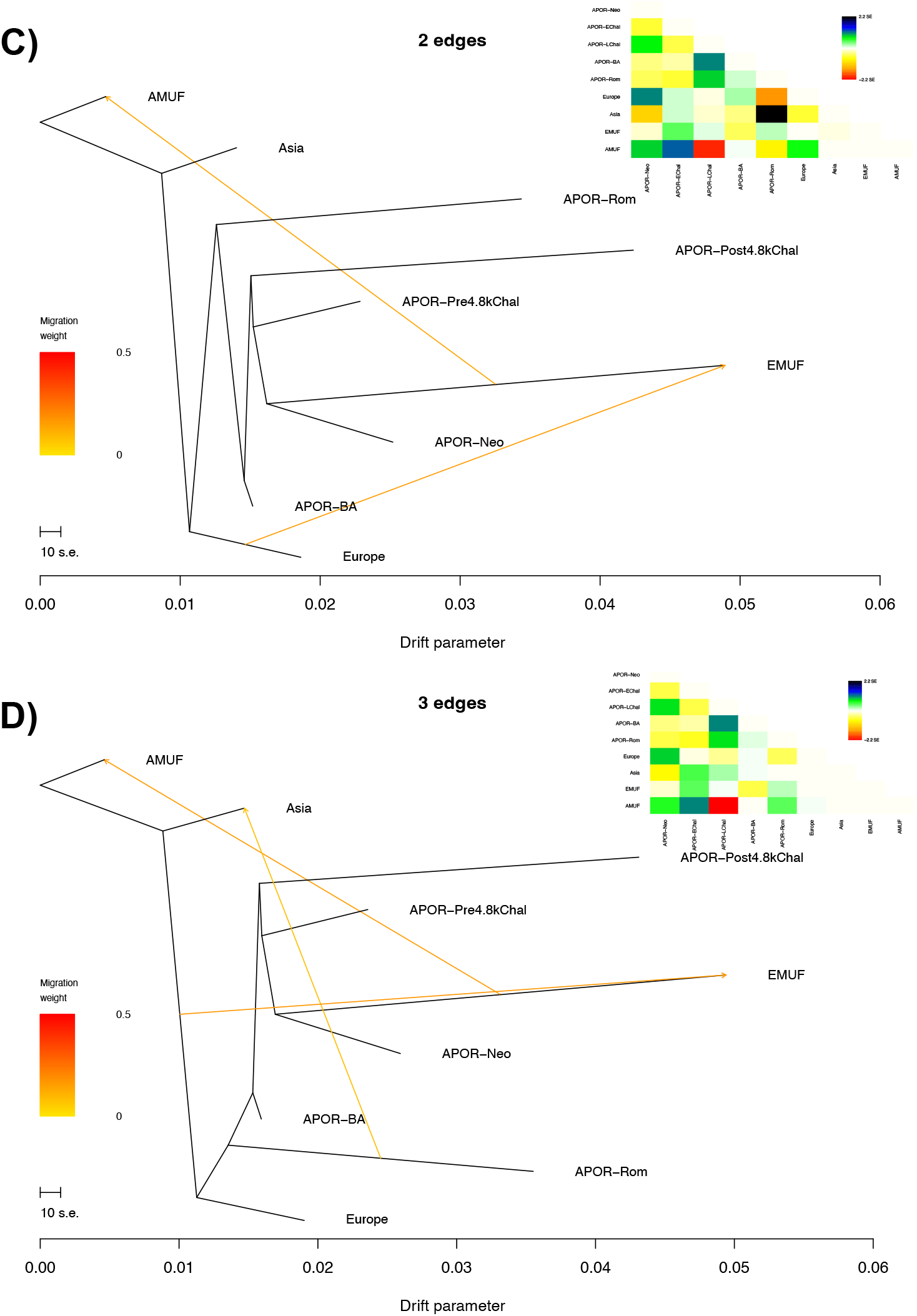
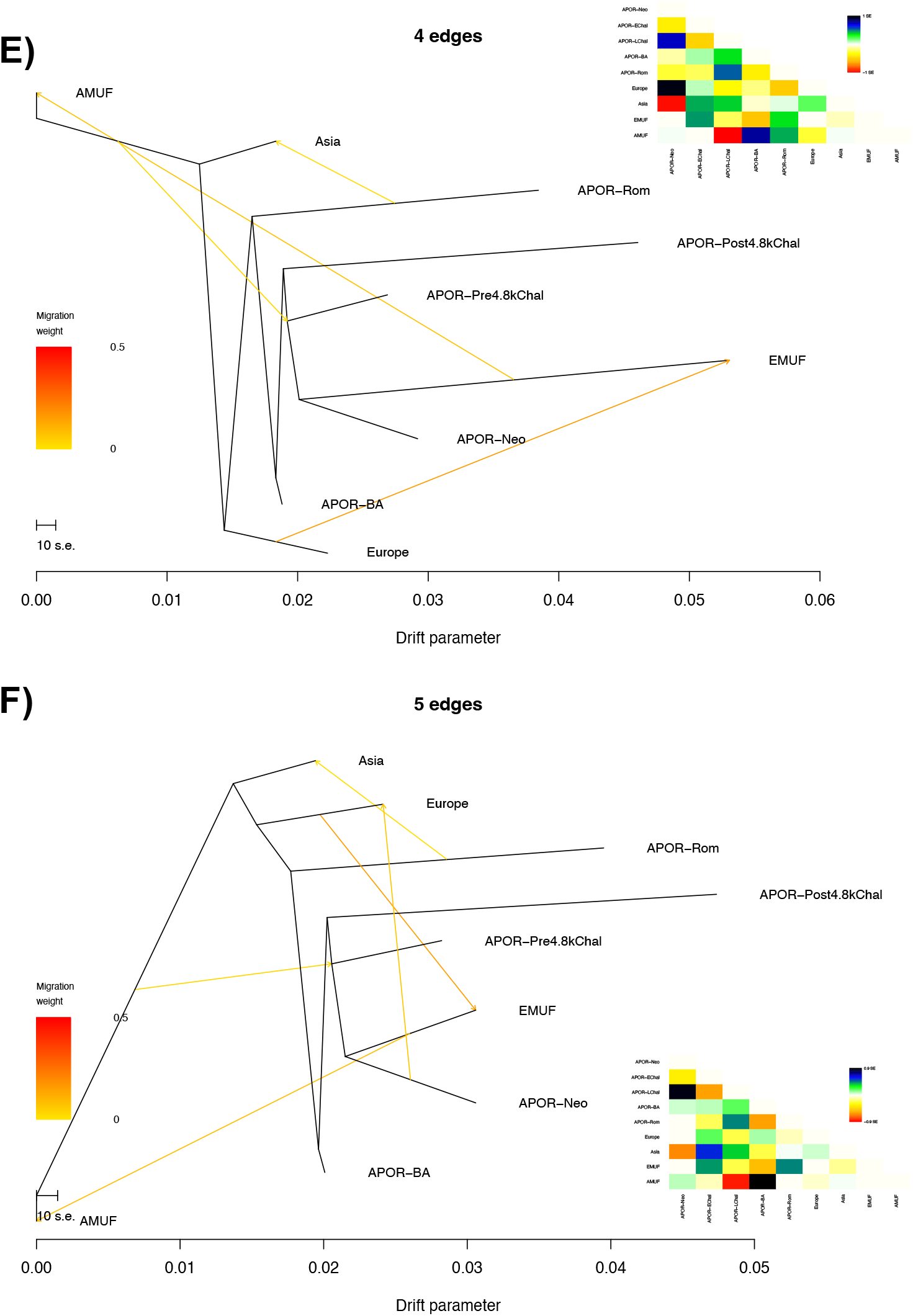
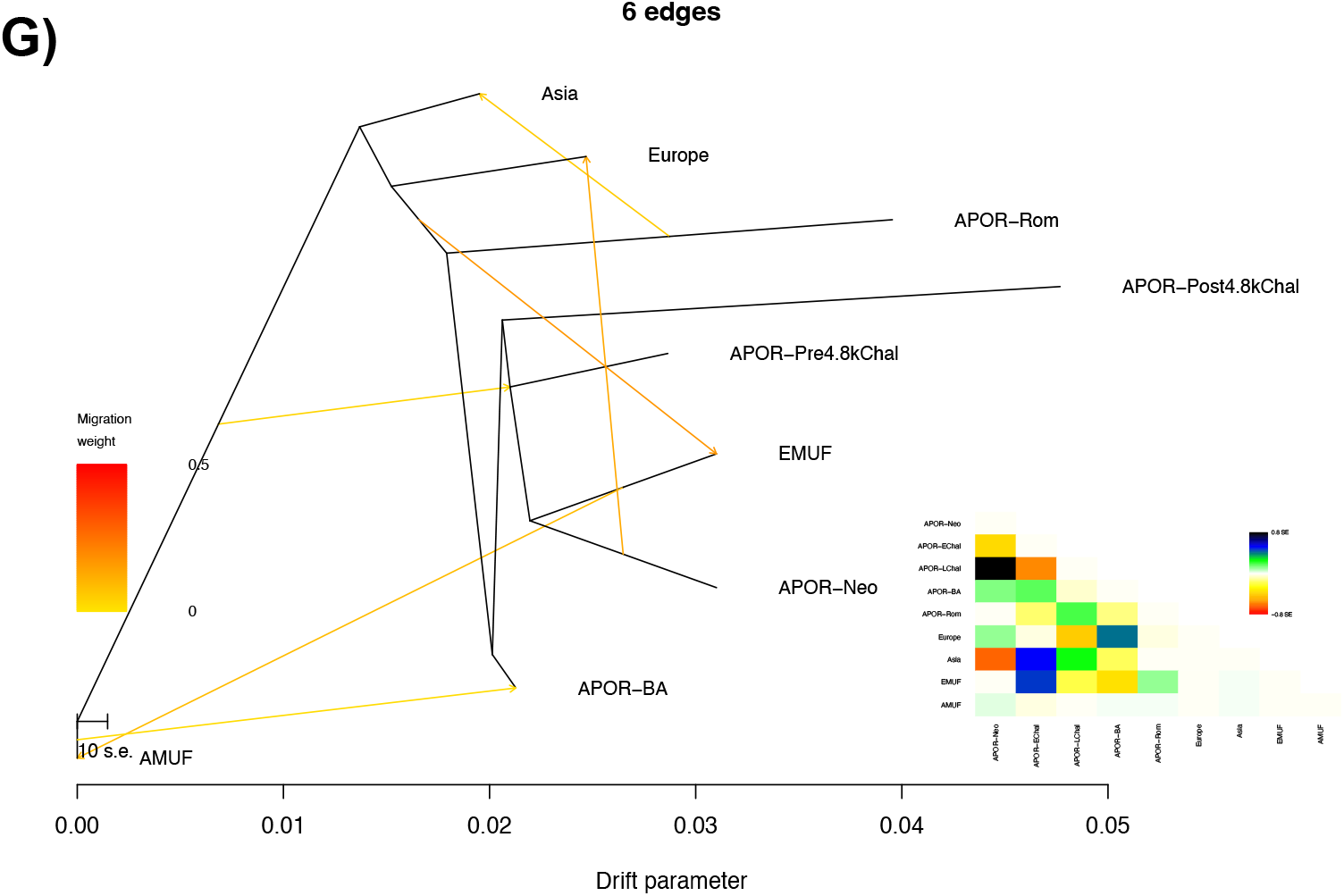
qpGraph admixture graphs allowing for 0 to 6 migration edges.

**Supplementary Figure 6.**
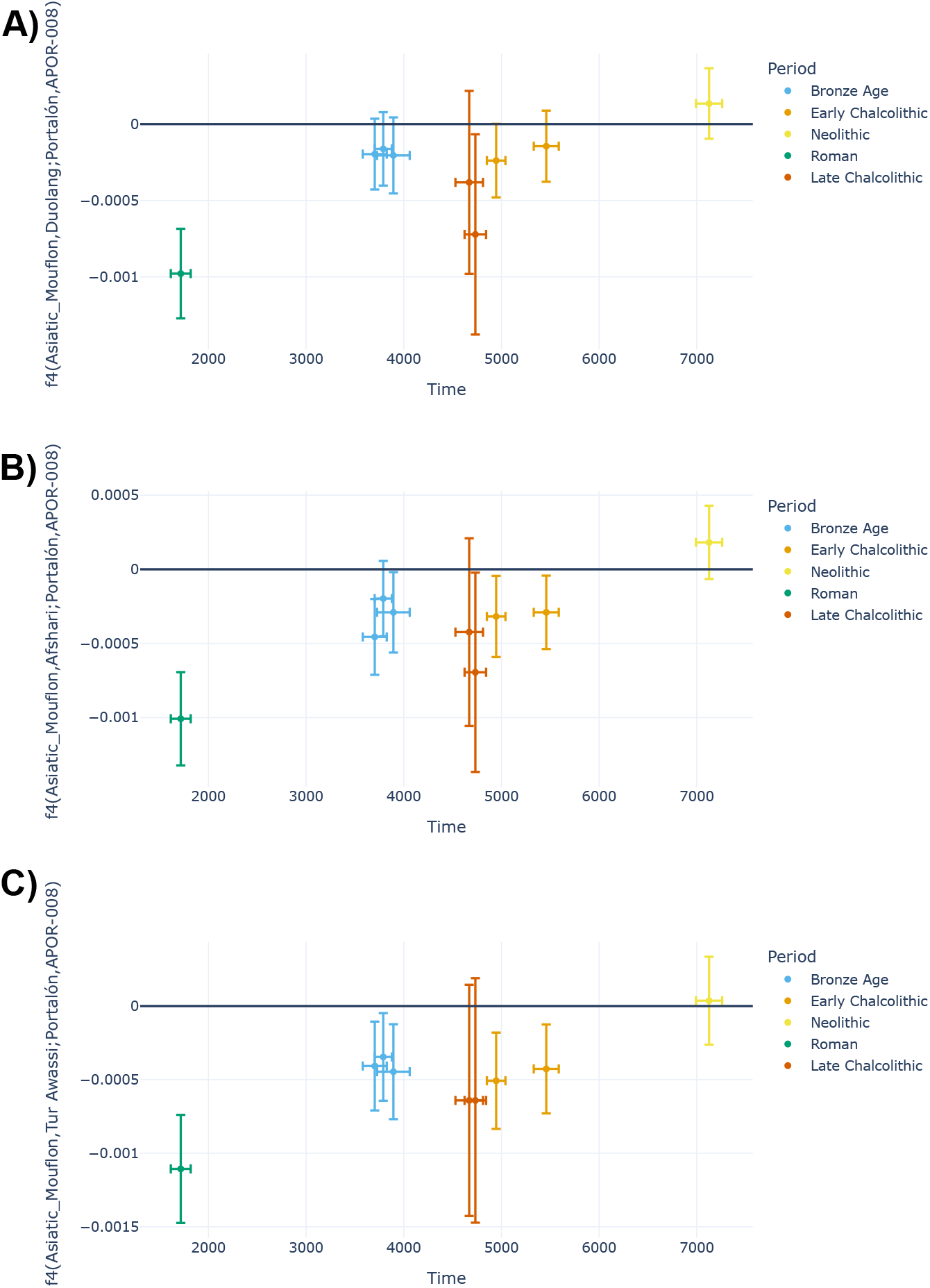
Values of the f_4_ statistics A) *f*_*4*_ *(Asiatic Mouflon, Duolang; El Portalón, El Portalón Neolithic)* B) *f*_*4*_ *(Asiatic Mouflon, Afshari; El Portalón, El Portalón Neolithic)* and C) *f*_*4*_ *(Asiatic Mouflon, Awassi; El Portalón, El Portalón Neolithic)* plotted against the radiocarbon dates of the ancient samples. “Early” and “Late” Chalcolithic refer to the Pre-4.8k and Post-4.8k Chalcolithic samples, respectively.

**Supplementary Figure 7.**
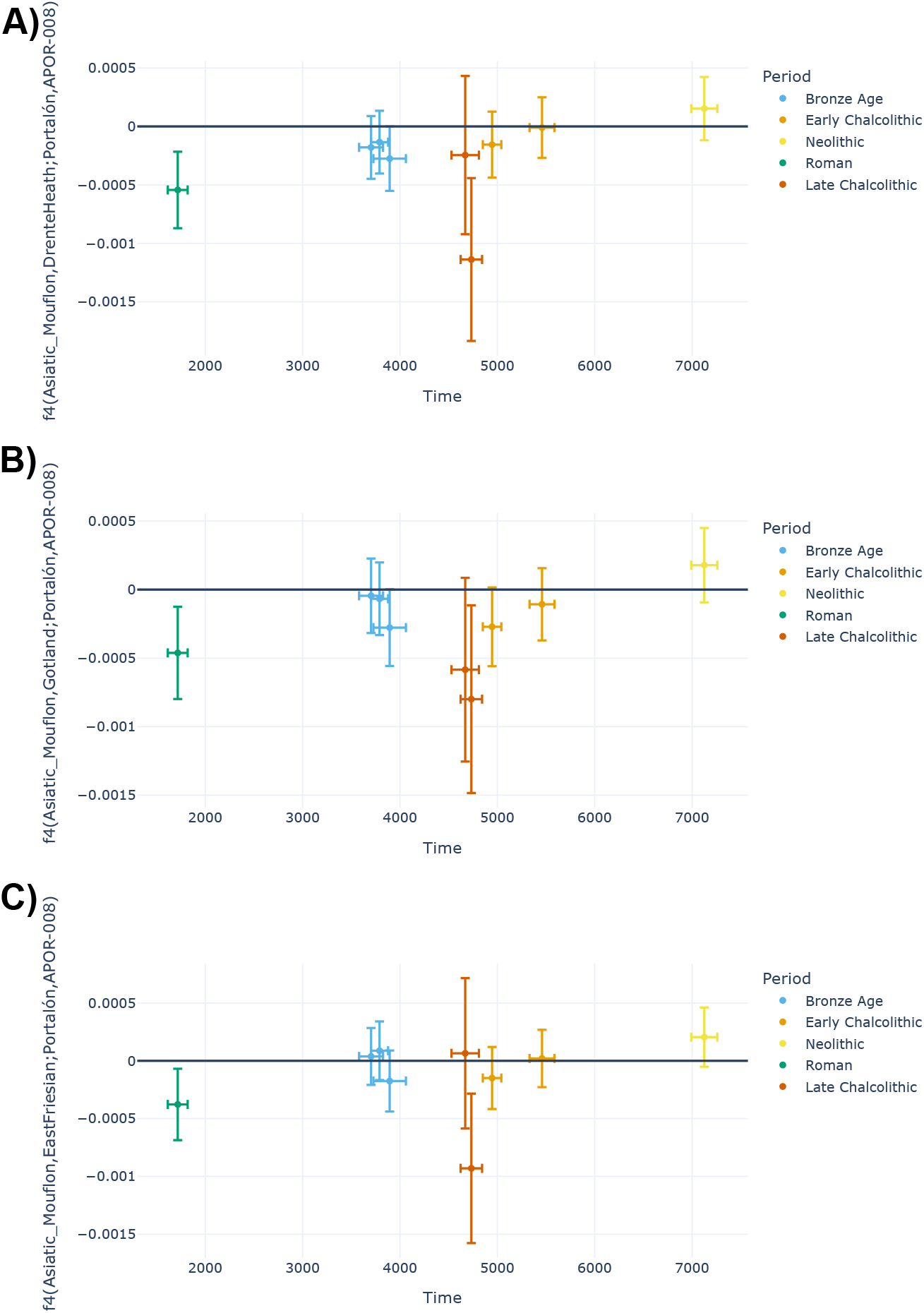
Values of the f_4_ statistics A) *f*_*4*_ *(Asiatic Mouflon, Drente Heathen; El Portalón, El Portalón Neolithic)*, B) *f*_*4*_ *(Asiatic Mouflon, Gotland; El Portalón, El Portalón Neolithic)* and C) *f*_*4*_ *(Asiatic Mouflon, East Friesian; El Portalón, El Portalón Neolithic)* plotted against the radiocarbon dates of the ancient samples. “Early” and “Late” Chalcolithic refer to the Pre-4.8k and Post-4.8k Chalcolithic samples, respectively.

## Notes

https://zenodo.org/records/18470345

### Competing Interest Statement

The authors have declared no competing interest.

